# RAB-35 aids apoptotic cell clearance by regulating cell corpse recognition and phagosome maturation

**DOI:** 10.1101/298380

**Authors:** Ryan C. Haley, Ying Wang, Zheng Zhou

**Affiliations:** Verna and Marrs McLean Department of Biochemistry and Molecular Biology, Baylor College of Medicine, Houston, TX 77030

## Abstract

In metazoans, apoptotic cells are swiftly engulfed by phagocytes and degraded inside phagosomes. Multiple small GTPases in the Rab family are known to function in phagosome maturation by regulating vesicle trafficking. We discovered *rab-35* as a new gene important for apoptotic cell clearance using an RNAi screen targeting putative Rab GTPases in *Caenorhabditis elegans*. We further identified TBC-10 as a putative GTPase-activating protein (GAP), and FLCN-1 and RME-4 as two putative Guanine Nucleotide Exchange Factors (GEFs), for RAB-35. RAB-35 function was found to be required for the incorporation of early endosomes to phagosomes and for the timely degradation of apoptotic cell corpses. More specifically, RAB-35 facilitates the switch of phagosomal membrane phosphatidylinositol species from PtdIns(4,5)P_2_ to PtdIns(3)P and promotes the recruitment of the small GTPase RAB-5 to phagosomal surfaces, processes that are essential for phagosome maturation. Interestingly, we observed that CED-1 performs these same functions, and to a much larger extent than RAB-35. Remarkably, in addition to cell corpse degradation, RAB-35 also facilitates the recognition of cell corpses independently of the CED-1 and CED-5 pathways. RAB-35 localizes to extending pseudopods and is further enriched on nascent phagosomes, consistent with its dual roles in regulating cell corpse-recognition and phagosome maturation. Epistasis analyses indicate that *rab-35* represents a novel third genetic pathway that acts in parallel to both of the canonical *ced-1/6/7* and *ced-2/5/10/12* engulfment pathways. We propose that RAB-35 acts as a robustness factor, leading a pathway that aids the canonical pathways for the engulfment and degradation of apoptotic cells.

## Introduction

During the development of metazoans, cells that undergo apoptosis are internalized and degraded by other cells that are referred to as engulfing cells or phagocytes (1–3). The phagocytic removal of apoptotic cells is an evolutionarily conserved event that supports normal tissue turnover and homeostasis, facilitates wound resolution and tissue regeneration, and prevents inflammatory and auto-immune responses induced by the release of dead cell contents (3,4). Throughout the development of *Caenorhabditis elegans* hermaphrodites, 300-500 of germ cells and 131 somatic cells undergo apoptosis (5–7). The temporal and spatial parameters of these cell death events are highly consistent between embryos (5). Apoptotic cells exhibit a “button-like” and highly refractive morphology under the Differential Interference Contrast (DIC) microscope, and are rapidly engulfed and degraded by multiple types of neighboring cells (5–8). Genetic screens and further characterizations of mutations that result in the “<u>*ce*</u>ll <u>*d*</u>eath abnormal” (Ced) phenotype, characterized by the accumulation of persistent cell corpses, have identified a number of genes that act in the recognition, engulfment, or degradation of cell corpses (9,10).

The Rab family of small GTPases play critical roles in membrane trafficking events, including endocytosis and exocytosis, autophagy, and phagosome maturation (11,12). A well-known mode of action is that Rab GTPases and their effectors serve as docking factors that facilitate the attachment and fusion of different membrane compartments and/or vesicles (11). Multiple mammalian and *C. elegans* Rab proteins play essential roles for phagosome maturation by facilitating the incorporation of intracellular organelles to phagosomes, an action that delivers digestive enzymes to the phagosomal lumen and that may also aid in the acidification of the lumen (13,14). *C. elegans* and mammalian RAB-5 are required for the recruitment and incorporation of early endosomes to phagosomes (15–17), while *C. elegans* and mammalian RAB-7 are critical for the incorporation of lysosomes to phagosomes (18,19). In *C. elegans*, both RAB-5 and RAB-7 function downstream of a signaling pathway that promotes phagosome maturation; this pathway is initiated by the phagocytic receptor CED-1 and mediated by the large GTPase DYN-1 (8,15,18,20). *C. elegans* RAB-2 and RAB-14 also make important contributions to the phagosomal degradation of cell corpses (21–23).

The signaling pathway led by CED-1 initiates phagosome maturation not only by recruiting Rab proteins to phagosomal surfaces, but also by initiating the production of PtdIns(3)P, a phosphorylated phosphatidylinositol species and an important second messenger, on phagosomal membranes (16,18). PtdIns(3)P recruits multiple effectors to phagosomes, including membrane remodeling factors and docking factors that facilitate the recruitment and fusion of intracellular vesicles (13,24). Consequently, phagosome maturation events are largely dependent on the presence of PtdIns(3)P and certain Rab GTPases (15,16). Interestingly, the presence of RAB-5 and PtdIns(3)P on phagosomal surfaces displays a co-dependent relationship (15,16).

Two PI3-kinases, PIKI-1 and VPS-34, catalyze the production of PtdIns(3)P on phagosomal surfaces (16,25). PIKI-1 and VPS-34 are functionally opposed by MTM-1, a PI3-phosphatase that dephosphorylates PtdIns(3)P and in this matter counteracts PI3-kinase activities (16). Throughout the phagosome maturation process, PtdIns(3)P is present on phagosomal surfaces in a two-wave oscillation pattern, a pattern coordinately regulated by PIKI-1, VPS-34, and MTM-1 (16). MTM-1 is recruited to the surface of extending pseudopods as an effector of PtdIns(4,5)P_2_, another phosphorylated phosphatidylinositol species that is enriched on the surface of growing pseudopods during engulfment (25). The initial appearance of PtdIns(3)P on phagosomes correlates not only with the recruitment of PIKI-1 to phagosomal surfaces by the CED-1 pathway (16), but also with the simultaneous disappearance of MTM-1 from nascent phagosomes, which is implicated to be a result of the disappearance of PtdIns(4,5)P_2_ from phagosomal surfaces (25). Whether the CED-1 signaling pathway also regulates the turnover of PtdIns(4,5)P_2_ has not yet been tested.

The phagocytic receptor CED-1 provides a link between the engulfment of apoptotic cells and the subsequent maturation of nascent phagosomes (8). During engulfment, CED-1 recognizes phosphatidylserine (PS), an “eat me” signal exposed on the surface of apoptotic cells, and defines one of the two canonical parallel pathways that stimulate pseudopod extension and cell corpse internalization (26–28). Several other key components act in this engulfment pathway alongside CED-1: CED-7, a homolog of mammalian ABC transporters that exposes PS on the surface of apoptotic cells; CED-6, an adaptor for CED-1; and DYN-1, an ortholog of the large GTPase dynamin that promotes “focal exocytosis” during pseudopod extension and stabilizes the cytoskeleton underneath extending pseudopods in response to CED-1 activation (8,29,30). In the other canonical engulfment pathway, CED-2 regulates the activity of the CED-5/CED-12 complex, presumably through its N-terminus that contains SH2 and SH3 domains (31,32). The CED-5/CED-12 complex, in turn, functions as a bipartite nucleotide exchange factor to activate the Rac GTPase CED-10 (33). CED-10 promotes the reorganization of the actin cytoskeleton and the extension of pseudopods around cell corpses (34,35). However, residual engulfment activity persists after inactivating both the *ced-1/-6/-7/dyn-1* and *ced-2*/*-5/-10*/*-12* pathways, suggesting that there are yet unknown pathways that play significant roles in cell-corpse engulfment (8,36). Although other proteins – such as alpha and beta integrins – are also reported to contribute to cell corpse engulfment, their effects are mild in comparison (37,38).

In addition to the putative missing pathways, many other ambiguities still surround the molecular mechanisms that control apoptotic cell clearance. For example, although many Rab GTPases have been implicated in the regulation of any clearance events, it remains unclear whether this is an exhaustive list. The *C. elegans* genome contains 30 genes that encode close homologs of mammalian Rab GTPases, 23 of which have been assigned names as *rab* genes (39). To determine which of these *rab* genes function in apoptotic cell clearance, we screened for any Rab GTPases that participate in cell corpse clearance using RNAi knockdown, excluding previously examined candidates such as *rab-2, −5, −7*, and *-14*. We discovered that inactivation of *rab-35*, which encodes a homolog of mammalian Rab35, caused a moderate yet significant Ced phenotype, indicating that RAB-35 functions in apoptotic cell clearance. Further characterizations revealed novel features and functions of RAB-35. Unlike RAB-5 and RAB-7, which are enriched on the surface of phagosomes and facilitate specific maturation events, RAB-35 weakly localizes at the extending pseudopods during engulfment, exhibits an ephemeral surge of localization on nascent phagosomes, and regulates multiple steps throughout apoptotic cell clearance. To facilitate the initiation of phagosome maturation, RAB-35 promotes the turnover of PtdIns(4,5)P_2_ and the recruitment of RAB-5, indirectly enabling phagosomal PtdIns(3)P production. Our findings further indicate that RAB-35 represents a clearance pathway that functions in parallel to the CED-1 and CED-5 pathways, yet in many ways resembles the mechanisms and functions of the CED-1 pathway. We thus propose that RAB-35 acts as a robustness factor and defines a new pathway that ensures the stability of apoptotic cell clearance.

## Results

### Inactivation of *rab-35* results in an increased number of persistent cell corpses

The *C. elegans* genome contains 30 genes that encode close homologs of mammalian Rab GTPases, 23 of which have been assigned gene names (40). Among these 23 putative *rab* genes, *rab-2*, *rab-5*, *rab-7*, and *rab-14* have been reported to act in the clearance of apoptotic cells (13). To determine if any other Rab proteins are involved in the same process, we individually knocked down the expression of 19 *rab* genes in *C. elegans* using the RNA interference (RNAi) treatment, and scored the number of germ cell corpses in the gonad of adult hermaphrodites (Materials and Methods). In addition, we scored the number of germ cell corpses in the *rab-10* deletion mutant (Materials and Methods). RNAi of *rab-1* and *rab-11.1* cause lethality before the worms develop into adults. Among the remaining 17 genes subject to RNAi treatment and the *rab-10(ok1494)* mutants, only 6 had more than four times the number of germ cell corpses compared to the wild-type control (Fig 1A). Of these 6, *rab-35*(RNAi) worms exhibited the highest number of persistent germ cell corpses, indicating the strongest defect in the clearance of germ cell corpses, characteristic of the Ced phenotype (Fig 1A). To verify this Ced phenotype, we examined two putative *rab-35* null alleles – the nonsense mutation *b1013* and the deletion allele *tm2058* (Fig 1B) (41). Both *rab-35(b1013)* and *rab-35(tm2058)* mutants exhibit identical Ced phenotypes in embryos in mid- (1.5-fold, ~420 min-post 1^st^ cleavage) and late- (late 4-fold, 700-800 min-post 1^st^ cleavage) stage embryos and the 48 hour post-L4 adult gonads (Fig. 1C-D), confirming the RNAi results.

**Fig 1.**
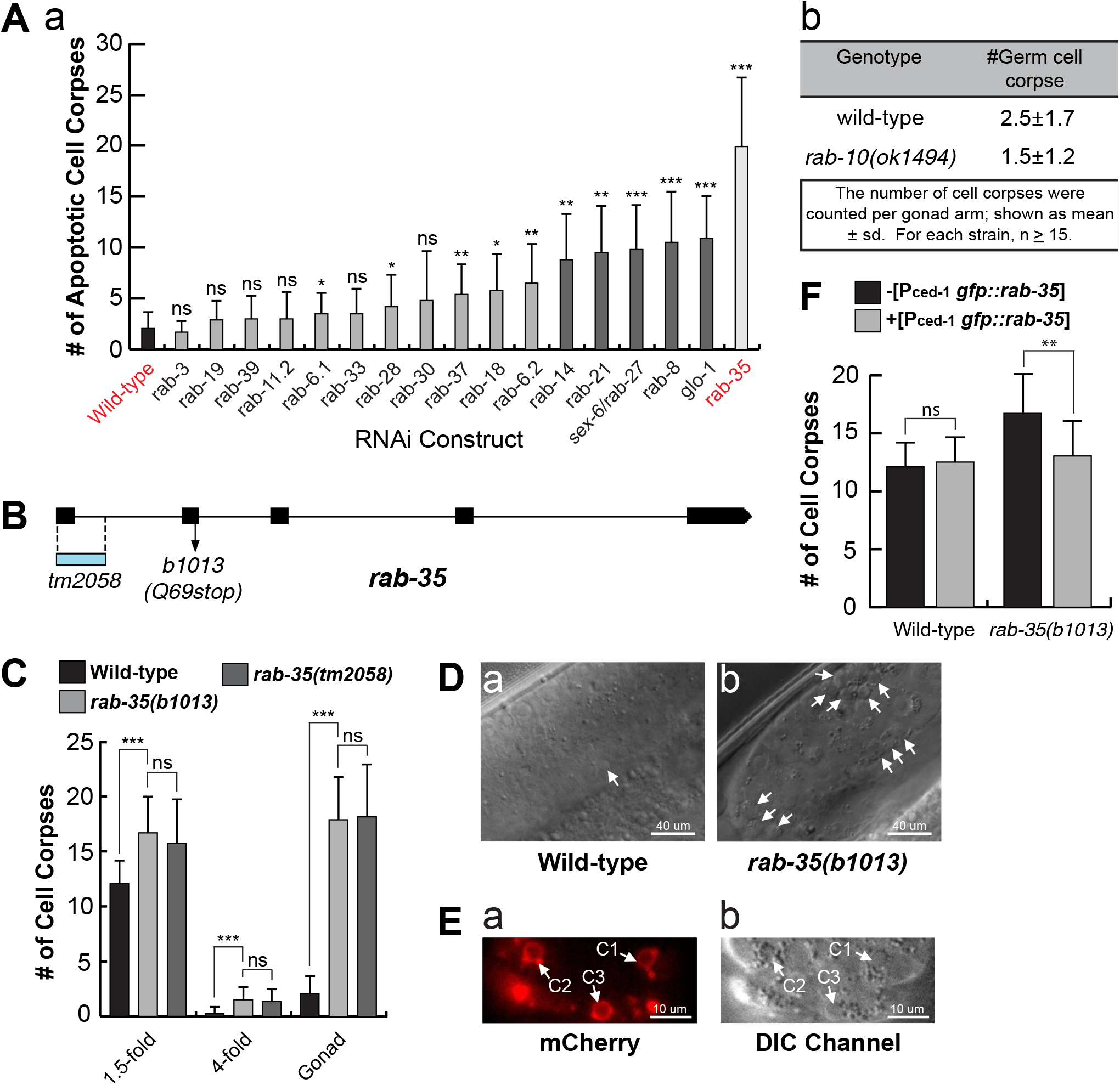
*rab-35(b1013)* mutants are defective in the clearance of apoptotic cells. All p-values were measured relative to wild-type. The student t-test was used for data analysis: *, 0.001 < p < 0.05; **, 0.00001 < p <0.001; ***, p <0.00001; ns, no significant difference. (A) The numbers of germ cell corpses were scored in 48-hour post-L4 adult gonads. A minimum of 15 animals were scored. Germ cell corpses were counted (a) after RNAi treatment of 17 *C. elegans* genes encoding RAB proteins, and (b) in wild-type and *rab-10(ok1494)* mutant strains. (B) The locations of the two null alleles of *rab-35* in the *rab-35* gene structure. Black rectangles mark exons. The blue rectangle indicates the location of the deletion in the *tm2058* allele. The arrow marks the position of the nonsense mutation carried in the *b1013* allele. (C) The numbers of cell corpses were scored at various developmental stages: 1.5-fold embryos, 4-fold embryos, and the 48-hour post-L4 adult gonad. For each data point, at least 15 animals were scored. Mean numbers were presented as bars. Error bars indicate standard deviation (sd). (D) Differential interference contrast (DIC) microscopy images of adult gonads of (a) wild-type and (b) *rab-35(b1013)* mutants. White arrows mark germ cell corpses. (E) The ventral surface of an *rab-35(b1013)* embryo that expresses MFG-E8::mcherry was visualized using both the mCherry (a) and DIC (b) channels at ~330 minutes post-first cleavage. White arrows mark the presence of MFG-E8::mcherry on C1, C2, and C3 in (a), and the cell corpses C1, C2, and C3 in (b). (F) The *gfp::rab-35* transgene expressed in engulfing cells is able to rescue the *rab-35* mutant phenotype. The mean numbers of cell corpses in 1.5-fold stage embryos in strains carrying or not carrying P_*ced-1*_ *gfp::rab-35* were presented in the bar graph. For each data point, at least 15 animals were scored. Error bars represent sd.

To determine whether the button-like objects observed under DIC optics in *rab-35(b1013)* mutants are actually cell corpses, we probed them for the exposure of PS on their surfaces, a distinct characteristic of cells undergoing apoptosis (27). Using MFG-E8::mCherry – a secreted PS-binding reporter (27), we detected bright mCherry signal specifically on the surface of the button-like objects (Fig 1E), indicating that they are indeed apoptotic cells. In addition, we expressed the *rab-35* cDNA, as an N-terminal GFP-tagged form, specifically in engulfing cells under the control of the *ced-1* promoter (P_*ced-1*_ *gfp::rab-35*) (26), and found that it completely rescued the Ced phenotype in *rab-35(b1013)* mutants (Fig 1F). This result suggests that the activity of RAB-35 in engulfing cells promotes cell corpse clearance.

RAB-35 is known to act in receptor-mediated endocytosis and endocytic recycling in *C. elegans* (41,42). We confirmed that the *rab-35* mutants have a characteristic excess of yolk in the pseudocoelom due to their inability to traffic yolk into oocytes as previously described (41,42) (Fig S1).

### RAB-35 localizes to developing pseudopods and must cycle between GDP- and GTP-bound states to function

Using time-lapse microscopy, we monitored the localization of GFP::RAB-35 expressed in engulfing cells (P_*ced-1*_*gfp::rab-35*), which rescues the Ced phenotype of *rab-35(b1013)* mutants. We tracked the clearance process of three apoptotic cells on the ventral surface: C1, engulfed by ABplaapppa; C2, engulfed by ABpraapppa; and C3, engulfed by ABplaapppp, using our previously established protocol (Fig 2A) (43). C1, C2, and C3 undergo apoptosis shortly after the initiation of ventral enclosure (~320-330 minutes post-first cleavage) (43). GFP::RAB-35 labels the extending pseudopods throughout engulfment; moreover, GFP::RAB-35 exhibits an ephemeral burst of enrichment on nascent phagosomes that lasts for 2-4 minutes (Fig 2B). Afterwards, the phagosomal GFP signal rapidly declines to the background level by approximately 15-20 minutes after the initiation of engulfment (Fig 2C). This dynamic enrichment pattern suggests that RAB-35 might participate in multiple events during apoptotic cell clearance.

**Fig 2.**
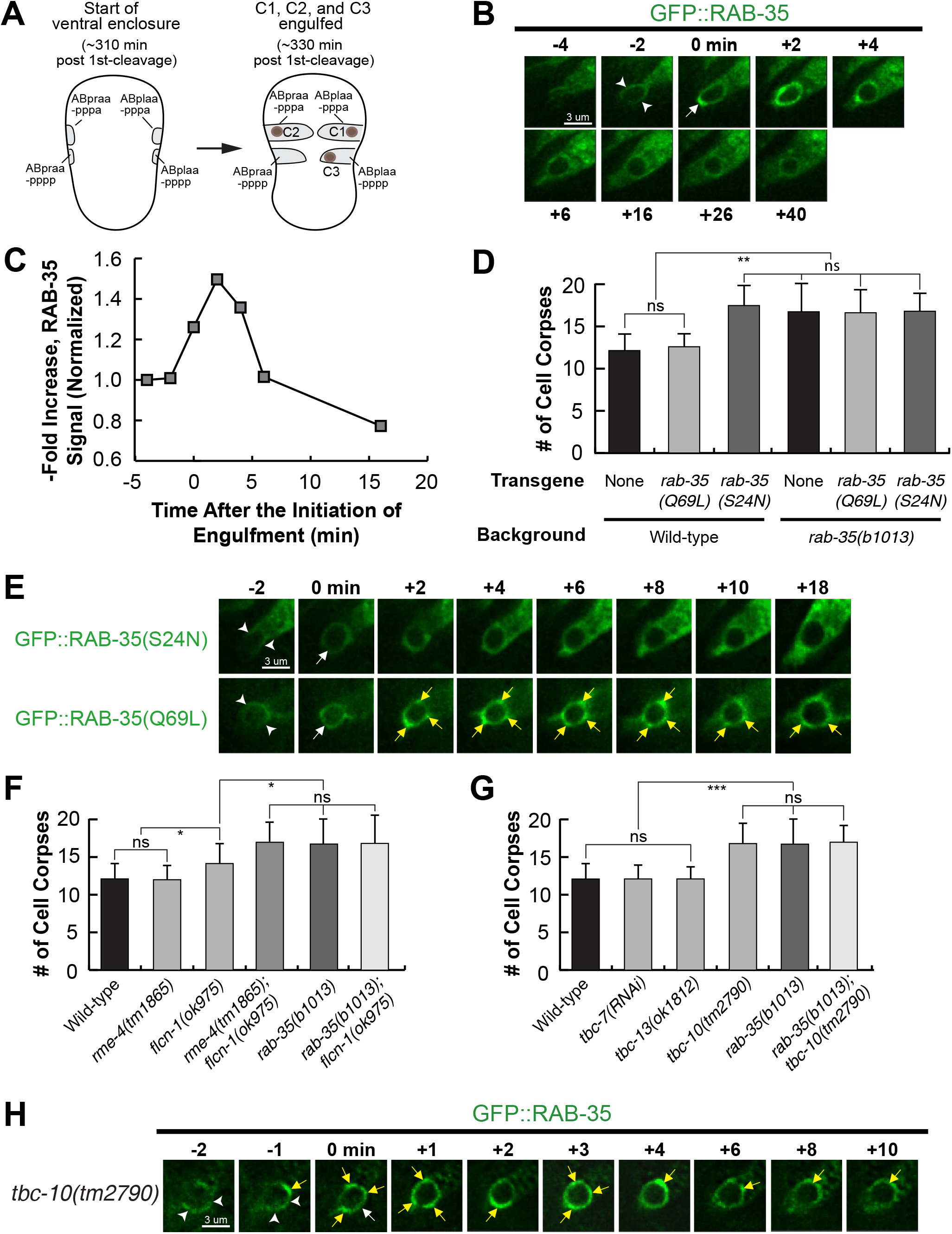
RAB-35 is localized to extending pseudopods and further enriched on nascent phagosomes. All GFP reporters are expressed in engulfing cells under the control of P_*ced-1*_. All p-values were measured relative to wild-type. The student t-test was used for data analysis: *, 0.001 < p < 0.05; **, 0.00001 < p <0.001; ***, p <0.00001; ns, no significant difference. (A) Diagram illustrating the features that allow for the visualization of ventral enclosure and apoptotic cell clearance. The start of ventral enclosure is defined as the moment the two ventral hypodermal cells (ABpraapppp and ABplaapppp) start extending to the ventral midline. Both the position of cell corpses C1, C2, and C3 (brown dots) as well as the identity of their engulfing cells are shown. (B) Time-lapse recording of GFP::RAB-35 during the engulfment and degradation of cell corpse C3 in a wild-type embryo. “0 min” indicates the formation of the nascent phagosome. Arrowheads mark the extending pseudopods. A whole arrow marks the nascent phagosome. (C) Graph showing the relative GFP::RAB-35 signal intensity over time on the surface of pseudopods and the phagosome in images shown in B. The fluorescence intensity of GFP was measured on the phagosomal surface and in the surrounding cytoplasm every 2 minutes, starting from the “0 min” time point. The phagosomal / cytoplasmic signal ratio over time was presented. Data is normalized relative to the signal ratio at the “0 min” time point. (D) The mean numbers of apoptotic cell corpses scored in 1.5-fold stage wild-type or *rab-35(b1013)* mutant embryos, in the presence or absence of transgenes overexpressing dominant negative GFP::RAB-35(S24N) or constitutively active GFP::RAB-35(Q69L), were presented in this bar graph. For each data point, at least 15 animals were scored. Error bars indicate sd. (E) The localization of GFP::RAB-35(S24N) and GFP::RAB-35(Q69L) during the engulfment of C3 and the early stage of phagosome maturation is presented in time-lapse images. “0 min” indicates the formation of the nascent phagosome. Arrowheads indicate extending pseudopods. A white arrow marks the nascent phagosome. Regions with enriched GFP::RAB-35(Q69L) signal on the phagosomal membrane are marked by yellow arrows. (F) Epistasis analysis between *rab-35* and genes that encode candidate GAP proteins for RAB-35. The mean numbers of apoptotic cell corpses scored in 1.5-fold stage wild-type and various single and double mutant combinations are presented in this bar graph. *tbc-7* was inactivated by RNAi. For each data point, at least 15 animals were scored. Error bars indicate sd. (G) Epistasis analysis between *rab-35* and genes that encode candidate GEF proteins for RAB-35. The mean numbers of apoptotic cell corpses scored in 1.5-fold stage wild-type and various single and double mutant combinations are presented in this bar graph. For each data point, at least 15 animals were scored. Error bars indicate sd. (H) Time-lapse recording of GFP::RAB-35 during the engulfment and degradation of cell corpse C3 in *tbc-10(tm2790)* mutant embryos. “0 min” indicates the formation of the nascent phagosome. Arrowheads mark the extending pseudopod. A white arrow marks the nascent phagosome. Regions with enriched GFP::RAB-35 signal on the phagosomal membrane are marked by yellow arrows.

We introduced S24N and Q69L, two point mutations previously established to convert Rab GTPases into the GDP-locked and GTP-locked forms (41), respectively, individually into the P_*ced-1*_ *gfp::rab-35* reporter constructs. Overexpression of RAB-35(S24N) produced a Ced phenotype in the wild-type background as strong as that displayed by *rab-35* null mutants (Fig 2D), verifying its predicted dominant-negative effect (41). Moreover, GFP::RAB-35(S24N) failed to enrich on the surfaces of extending pseudopods or nascent phagosomes (Fig 2E), suggesting that it is a non-functional form. Remarkably, overexpression of RAB-35(Q69L), the presumed GTP-locked form, failed to rescue the Ced phenotype of *rab-35* mutants (Fig 2D), although it displayed persistent enrichment on the phagosomal membrane (Fig 2E). Together, the altered localization patterns and the lack of rescuing activity observed in both mutant forms of RAB-35 suggest that the cycling of RAB-35 between the GDP-bound and GTP-bound states is required for its proper localization and function during apoptotic cell clearance.

### Determining the putative GAP and GEFs for RAB-35 for apoptotic cell clearance

To better understand how the cycling of RAB-35 between the GDP- and GTP-bound forms is regulated, we examined *C. elegans* orthologs of known GAPs and GEFs of mammalian Rab35 to determine which ones function in the context of apoptotic cell clearance. We first studied the loss-of-function alleles of *rme-4* and *flcn-1*, which encode the *C. elegans* orthologs of the mammalian GEFs connecdenns 1/2/3 and folliculin, respectively (44). The *flcn-1(ok975)* null mutation resulted in a Ced phenotype that is slightly weaker than that of *rab-35(b1013)* mutants (Fig 2F). Furthermore, the *flcn-1(ok975)*; *rab-35(b1013)* double mutants exhibited a Ced phenotype identical to that of *rab-35(b1013)* mutants, placing both *flcn-1* and *rab-35* in the same genetic pathway (Fig 2F). These results suggest that *flcn-1* might act as a GEF for RAB-35, but also that it may be working in tandem with another GEF.

RME-4 was previously reported to act as a GEF for RAB-35 during its function in endocytic trafficking (41). Interestingly, although *rme-4(tm1865)* single mutants did not exhibit any statistically significant Ced phenotype, the *flcn-1(ok975)*; *rme-4(tm1865)* double mutants exhibited a Ced phenotype more severe than that displayed by the *flcn-1(ok975)* single mutants and identical in severity to that of *rab-35(b1013)* mutants (Fig 2F), suggesting that RME-4 might also function as a GEF for RAB-35 in the context of apoptotic cell clearance. Compared to FLCN-1, however, the contribution of RME-4 to apoptotic cell clearance is relatively minor and redundant.

We subsequently probed deletion mutant alleles of the genes *tbc-7*, *tbc-10*, and *tbc-13* (www.wormbase.org), which encode the *C. elegans* orthologs of TBC1D24, TBC1D10A/B/C, and TBC1D13, known GAPs for mammalian Rab35, respectively (41) (Fig S2). We found considerable evidence that TBC-10 acts as the sole GAP for RAB-35 in the context of cell corpse clearance. Firstly, a loss-of-function mutant of *tbc-10*, but not those of *tbc-7* or *tbc-13*, exhibits a Ced phenotype identical to that of *rab-35(b1013)* mutants (Fig 2G). These results are also consistent with our previous observation that RAB-35 must cycle between its GTP- and GDP-bound forms to function, as inactivating its putative GAP (*tbc-10*) also appears to disable RAB-35. If the GTP-locked form of RAB-35 was its active form, *tbc-10* mutants would instead lock RAB-35 in a constitutively active form and thus fail to exhibit a Ced phenotype. Secondly, when we tracked the localization of GFP::RAB-35 throughout the clearance of C1, C2, and C3 in *tbc-10* mutants, we found that – relative to wild-type – RAB-35 localized to extending pseudopods and nascent phagosomes normally, but that its removal from phagosomal surfaces was delayed (Fig 2H). This pattern is similar to that of GFP::RAB-35(Q69L) in a wild-type background (Fig 2E), indicating that GFP::RAB-35 is locked in the GTP-bound form in *tbc-10* mutants. Finally, the *tbc-10(tm2790); rab-35(b1013)* double mutants did not enhance the Ced phenotype over either single mutant, confirming that *tbc-10* is in the same genetic pathway as *rab-35*, as would be expected for a putative GAP for RAB-35 (Fig 2G).

### *rab-35* loss of function causes delays in phagosomal maturation

*C. elegans* RAB-2, RAB-5, and RAB-7 all play important roles in the maturation of phagosomes that contain apoptotic cells (13). To determine whether RAB-35 is involved in this process, we measured how fast phagosomes degraded the cell corpses C1, C2, or C3 in *rab-35* mutant embryos. The lifetime of a phagosome is measured using a combination of a GFP::moesin reporter, which specifically labels the polymerized actin filaments underneath the extending pseudopods (28), and CTNS-1::mRFP, a lysosomal membrane marker that is enriched on the surface of phagosomes during maturation (18). These two reporters are co-expressed in engulfing cells under the control of the P_*ced-1*_ promoter (18,28). GFP::moesin is used to determine when pseudopods fuse to form a nascent phagosome, providing a way to determine the time point when phagosomal maturation begins and to measure the initial diameter of a nascent phagosome. CTNS-1::mRFP is then used to track and measure the diameter of a phagosome throughout maturation (Fig 3A). We defined phagosomal lifetime as how long it takes for the phagosome to shrink to one-third of its original radius after the initiation of phagosome maturation. Using this assay, we found that *rab-35(b1013)* mutants exhibited a significantly longer phagosomal lifetime than their wild-type counterparts; 75% of phagosomes in *rab-35* mutants had a lifetime longer than 60 minutes, compared to only 13.3% of phagosomes in wild-type embryos (Fig 3B). These results indicate that RAB-35 is important for the efficient degradation of phagosomal contents.

**Fig 3.**
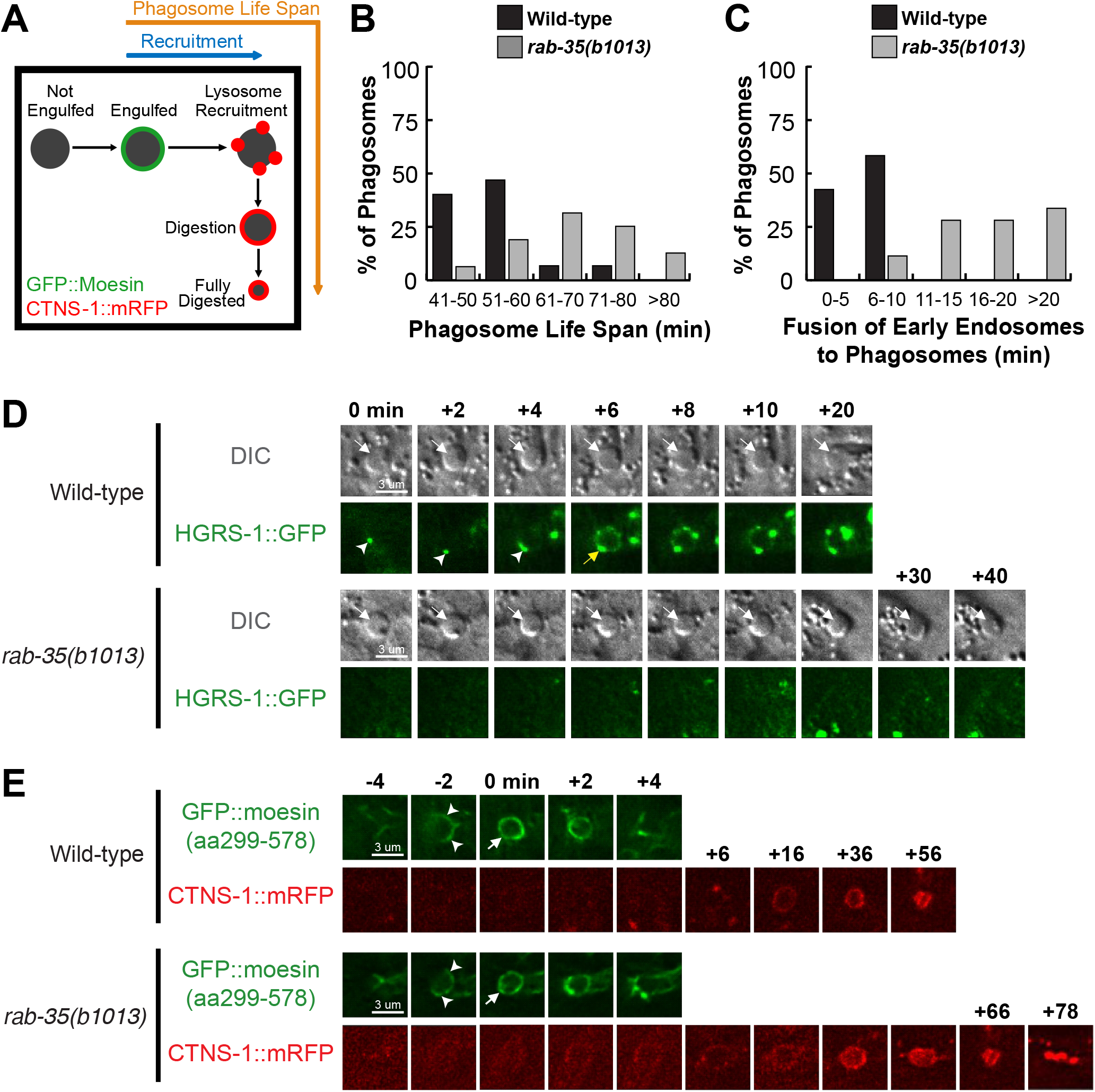
*rab-35* mutants exhibit delays in the recruitment of early endosomes, but not lysosomes, to phagosomes. (A) Diagram outlining the experiment strategy to measure the life span of a phagosome. GFP::moesin(aa299-578), which labels pseudopods, serves to mark the “0 min” time point of the formation of a nascent phagosome, while CTNS-1::mRFP, a lysosome marker, acts to track the recruitment and fusion of lysosomes to the phagosome as well as to label the phagosome during the subsequent digestion of the cell corpse. (B) Histogram displaying the life span of phagosomes bearing cell corpses C1, C2, and C3 in wild-type and *rab-35(b1013)* embryos. The life span of a phagosome is defined as the time interval between the “0 min” time point when a nascent phagosome is initially formed and the time point when a phagosome shrinks to one-third of its measured radius at “0 min”. For each genotype, at least 15 phagosomes were scored. (C) Histogram displaying the range of time it takes for early endosomes to be recruited to the phagosomal surface in wild-type and *rab-35(b1013)* embryos. Phagosomes bearing cell corpses C1, C2, and C3 were scored. The time span of early endosome recruitment is measured as the time interval between “0 min” and the time point when the accumulating early endosomes first form a continuous ring around a phagosome. For each genotype, at least 15 phagosomes were scored. (D) Time-lapse images monitoring the recruitment of early endosomes (reporter: HGRS-1::GFP) to the phagosomal surface after a phagosome forms (the “0 min” time point). The monitored cell corpses (white arrows) are visualized using DIC microscopic images. Arrowheads indicate extending pseudopods. The GFP ring, when it is first completed around the phagosome, is labeled with a yellow arrow. (E) Time-lapse image series showing the processes of the engulfment and the degradation process of a phagosome bearing the cell corpse C3 in each of the wild-type and *rab-35(b1013)* embryos, using GFP::moesin(aa299-578) as the pseudopod reporter and CTNS-1::mRFP as a lysosome marker. “0 min” indicates the formation of the nascent phagosome. Arrowheads indicate extending pseudopods. A white arrow marks the nascent phagosome.

### *rab-35* mutants are defective in the incorporation of early endosomes to phagosomes

During the degradation of cell corpses, two kinds of intracellular organelles – early endosomes and lysosomes – are recruited to the surface of phagosomes and subsequently fuse to the phagosomal membrane, depositing their contents into the phagosomal lumen (13). The recruitment and fusion of early endosomes to the phagosome was probed using HGRS-1::GFP, an established early endosomal surface marker expressed in engulfing cells (8). Starting from the birth of a nascent phagosome, HGRS-1::GFP appears on the surface of a phagosome as puncta (Fig 3D). The continuous accumulation of the puncta over time generates a GFP ring around the phagosomal surface (Fig 3D). In *rab-35* mutant embryos, this GFP ring appears much slower relative to wild-type embryos, suggesting that RAB-35 function is required for the efficient recruitment of early endosomes (Fig 3C-D).

Likewise, CTNS-1::mRFP was used to visualize the recruitment of lysosomes to the surface of phagosomes. CTNS-1::mRFP first appeared on phagosomal surfaces as puncta, with the accumulating puncta gradually forming a mRFP ring on the phagosomal surface over time (18). Time-lapse recording and quantitative analysis found that *rab-35(b1013)* mutants had no statistically significant delays in the recruitment of lysosomes (Fig S3B). To further determine if the fusion of lysosomes to phagosomes was normal in *rab-35* mutants, we monitored the entry of a NUC-1::mRFP reporter (expressed in engulfing cells under P_*ced-1*_) into the phagosomal lumen. NUC-1 is an endonuclease that specifically resides in the lysosomal lumen (45). Similar to CTNS-1::mRFP, NUC-1::mRFP is recruited to phagosomal surfaces as mRFP^+^ puncta (45); however, unlike CTNS-1, the fusion of lysosomes to the phagosome causes NUC-1::mRFP to enter the phagosomal lumen (Fig S3A). In *rab-35(b1013)* embryos, the entry of the NUC-1::mRFP signal occurred on an timescale identical to that of wild-type embryos, suggesting that inactivating *rab-35* has no effect on phagolysosomal fusion (Fig S3C).

### RAB-35 promotes the initiation of phagosomal maturation by facilitating the PtdIns(4,5)P_2_ to PtdIns(3)P shift

The rapid enrichment of RAB-35 during the formation of a nascent phagosome (Fig 2B) suggests that RAB-35 may function during the initiation of phagosome maturation. At this same time, the predominant phosphatidylinositol species on the phagosomal surface switches from PtdIns(4,5)P_2_ to PtdIns(3)P, a process necessary for the progression of phagosome maturation and cell corpse degradation (16,18,28). We monitored the dynamic localization pattern of RAB-35 relative to those patterns of PtdIns(4,5)P_2_ and PtdIns(3)P. This was done by co-expressing our own RAB-35 reporters with previously established reporters for PtdIns(4,5)P_2_ (PH::GFP) or PtdIns(3)P (2xFYVE::mRFP) (18,28) (Fig 4A-B). We observed that RAB-35 enrichment corresponded exactly with both the loss of PtdIns(4,5)P_2_ and the gain of PtdIns(3)P on the surface of a nascent phagosome (Fig 4A-B).

**Fig 4.**
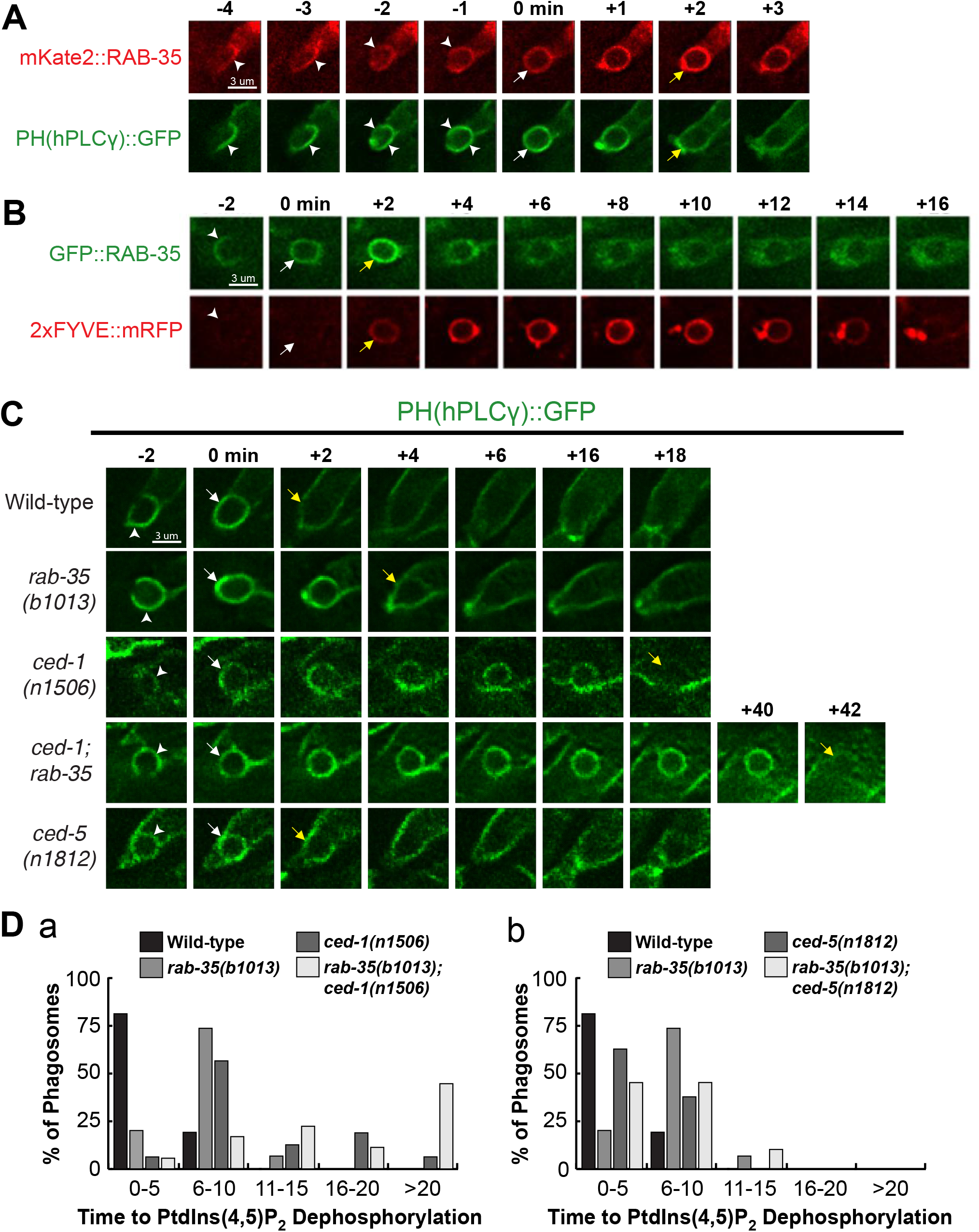
RAB-35 is enriched on phagosomal surfaces during the PtdIns(4,5)P_2_ to PtdIns(3)P shift and functions in PtdIns(4,5)P_2_ removal. (A) Time-lapse images during and after the formation of a phagosome carrying C3 in a wild-type embryo co-expressing P_*ced-1*_ mKate2::rab-35 and the PtdIns(4,5)P_2_ marker P_*ced-1*_ *PH(hPLCγ)::gfp*. “0 min” indicates the formation of the nascent phagosome. Arrowheads indicate extending pseudopods. White arrows mark the nascent phagosome, while yellow arrows mark both the gain of mKate2::RAB-35 and the loss of PH(hPLCγ)::GFP from the phagosomal surface. (B) Time-lapse images during and after the formation of a phagosome carrying C3 in a wild-type embryo co-expressing P_*ced-1*_ *gfp::rab-35* and the PtdIns(3)P marker P_*ced-1*_ 2xFYVE::mRFP. “0 min” indicates the formation of the nascent phagosome. Arrowheads indicate extending pseudopods. White arrows mark the nascent phagosome, while yellow arrows mark both the gain of GFP::RAB-35 and the loss of 2xFYVE::mRFP. (C) Time-lapse images during and after the formation of a phagosome carrying C3 in embryos of different genotypes expressing P_*ced-1*_ *PH(hPLCγ)::gfp*. “0 min” indicates the formation of the nascent phagosome. Arrowheads indicate extending pseudopods. White arrows mark the nascent phagosome, while yellow arrows mark the first time point when PtdIns(4,5)P_2_ is no longer observed on the phagosome surface. (D) Histograms displaying the range of time it takes for the disappearance of PtdIns(4,5)P_2_ from the surface of phagosomes bearing C1, C2, and C3 in embryos of various genotypes. *rab-35* mutants are coupled with null mutations in *ced-1* (a) and *ced-5* (b). The time span of PtdIns(4,5)P_2_ disappearance is scored as the time interval between the formation of a nascent phagosome (“0 min”) and the first time point)::GFP signal is no longer enriched on the phagosomal surface. For each genotype, at least 15 phagosomes were scored.

Because PtdIns(3)P is essential for phagosome maturation (see Introduction), and because the disappearance of PtdIns(4,5)P_2_ from phagosomal surfaces is correlated with the production of PtdIns(3)P on phagosomes (25), we examined whether RAB-35 regulates the dynamic pattern of PtdIns(4,5)P_2_ and PtdIns(3)P on phagosomes. We first monitored PtdIns(4,5)P_2_ dynamics on the surface of phagosomes in a series of mutant embryos (Fig 4C). We found that in *rab-35(b1013)* and *ced-1(n1506)* mutants, but not *ced-5(n1812)* mutants, PtdIns(4,5)P_2_ persists longer on phagosomal surfaces (Fig 4C-D). *ced-1(n1506)* mutants exhibited a longer delay in PtdIns(4,5)P_2_ disappearance than *rab-35(b1013)* mutants, and *rab-35(b1013); ced-1(n1506)* double mutants exhibited a much more severe delay than either single mutant (Fig 4D(a)). These results suggest that RAB-35 and CED-1 act in a partially redundant fashion for the removal of PtdIns(4,5)P_2_ from phagosomal surfaces.

The phagocytic receptor CED-1 was previously demonstrated to play an essential role in initiating PtdIns(3)P synthesis on the surface of nascent phagosomes (18). In *rab-35* mutant embryos, the appearance of the initial peak of PtdIns(3)P was significantly delayed relative to wild-type, although the defect was not as strong as that observed in *ced-1* mutant embryos (Fig 5A+B(a)). *rab-35; ced-1* double mutants have a stronger delay in the PtdIns(3)P appearance compared to either single mutant (Fig 5B(a)), again suggesting that RAB-35 and CED-1 act in a partially redundant fashion for the generation of PtdIns(3)P on phagosomal surfaces. However, because *ced-5* mutants fail to exhibit any delay in PtdIns(3)P production, and because the severity of the delay of PtdIns(3)P displayed by the *rab-35; ced-5* double mutants is equivalent to that of the *rab-35* single mutants, we conclude that *ced-5* is not involved in the regulation of PtdIns(3)P production (Fig 5A+B(b)).

**Fig 5.**
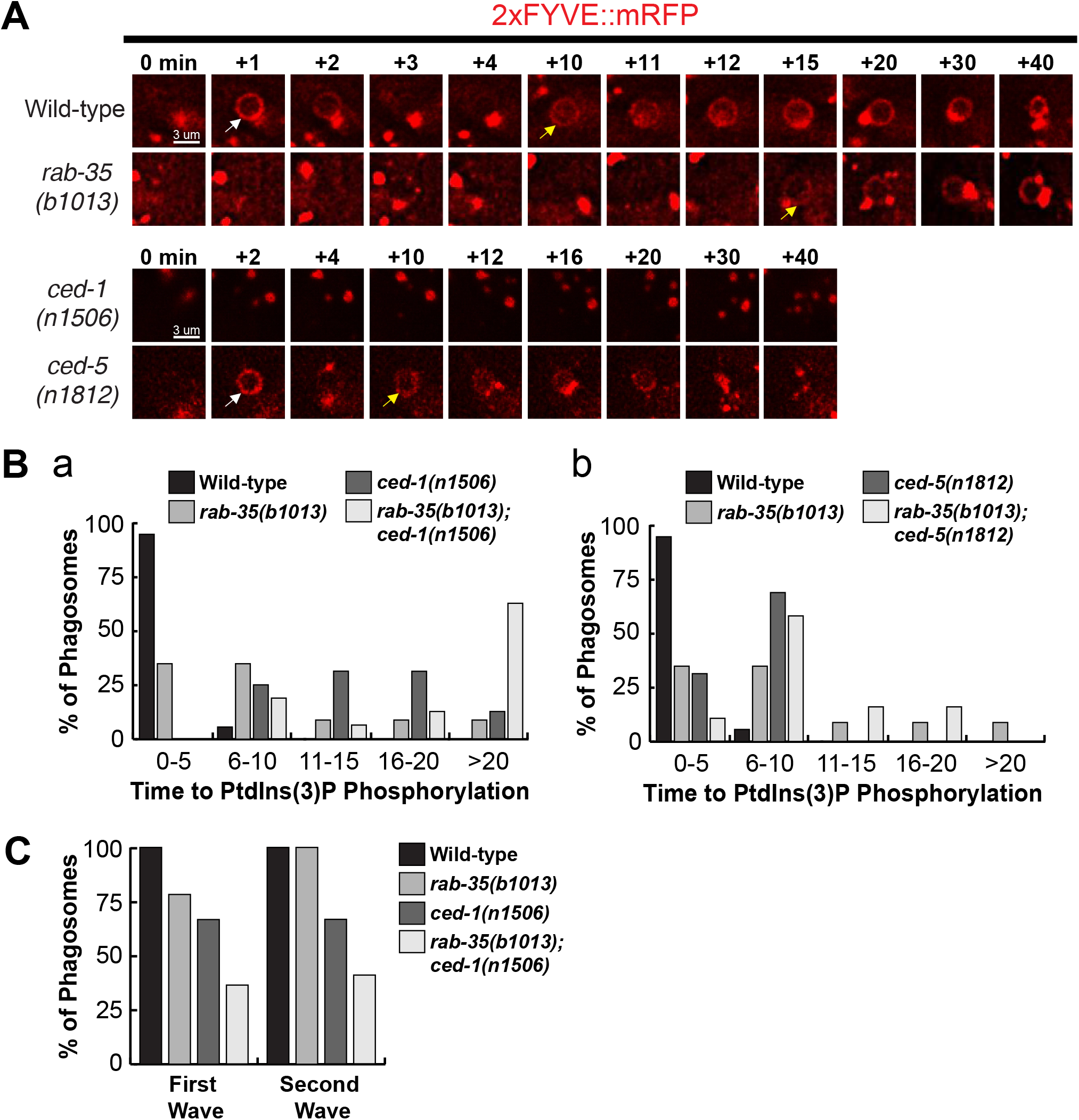
*rab-35* and *ced-1* function in parallel to produce PtdIns(3)P on the phagosomal membrane. (A) Time-lapse images during and after the formation of a phagosome carrying C3 in embryos of different genotypes expressing P_*ced-1*_*2xFYVE::mRFP*. “0 min” indicates the formation of the nascent phagosome, determined using the pseudopod marker GFP::moesin(aa299-578) [*not shown*]. White arrows indicate the time point when 1^st^ wave of PtdIns(3)P appears on the nascent phagosome. Yellow arrows mark the time point when the 2^nd^ wave of PtdIns(3)P appears on the phagosome. (B) Histogram displaying the range of time it takes for the 1^st^ peak of PtdIns(3)P to appear on phagosomes in wild-type, *rab-35(b1013)*, and either *ced-1(n1506)* and *rab-35(b1013)*; *ced-1(n1506)* (a) or *ced-5(n1812)* and *rab-35(b1013)*; *ced-5(n1812)* (b) embryos. Phagosomes bearing cell corpses C1, C2, and C3 were scored. “0 min” indicates the formation of the nascent phagosome. This time interval is defined as that between the formation of a nascent phagosome (“0 min”) and the first time point that the 1^st^ wave of PtdIns(3)P appears on the phagosome surface. For each genotype, at least 15 phagosomes were scored. (C) The frequency of appearance of the 1^st^ and 2^nd^ peaks of PtdIns(3)P on phagosomes carrying C1, C2, and C3 in wild-type, *rab-35(b1013)*, *ced-1(n1506)*, and *rab-35(b1013); ced-1(n1506)* embryos. For each genotype, at least 15 phagosomes were scored.

PtdIns(3)P typically appears on the phagosomal surface in two distinct waves (16), as exhibited in Figure 5A (time-lapse strip of a wild-type embryo, white and yellow arrows). We thus quantified whether the first and second waves of PtdIns(3)P were present in each of the aforementioned mutants. In stark contrast to wild-type embryos, where every phagosome exhibited both PtdIns(3)P waves, 21.7% of phagosomes in *rab-35(b1013)* mutants failed to produce the first wave of PtdIns(3)P, while 33.3% of phagosomes in *ced-1(n1506)* mutants failed to produce either the first or second wave (Fig 5C). *rab-35(b1013)*; *ced-1(n1506)* double mutants exhibited more severe defects than either single mutant (Fig 5C). Together, our observations indicate that RAB-35 and CED-1 act in parallel to promote the production of PtdIns(3)P on the surface of nascent phagosomes.

### RAB-35 is required for the efficient removal of MTM-1 from phagosomal surfaces

To further explore how loss of function of *rab-35* delays PtdIns(3)P production, we characterized the localization of PtdIns(3)P kinases and phosphatases using time-lapse recording of phagosomes containing C1, C2, and C3. Although both PIKI-1 and VPS-34 function as PI3-kinases during phagosome maturation, only PIKI-1 is observed when the nascent phagosome seals and initiates the PtdIns(4,5)P_2_ to PtdIns(3)P shift, while VPS-34 functions later on in maturation (16). Therefore, we monitored the phagosomal dynamics of PIKI-1, as well as MTM-1, the PI3-phosphatase it largely antagonizes (16,25). *rab-35(b1013)* mutants exhibited normal recruitment of PIKI-1::GFP; the enrichment of PIKI-1::GFP was observed on the surface of every phagosome, and the level of enrichment was comparable to that observed in wild-type embryos (Fig S4). In contrast, MTM-1::GFP persists on the surface of phagosomes approximately twice as long in *rab-35(b1013)* mutants – and more than thrice as long in *ced-1(n1056)* mutants – compared to wild-type embryos (Fig 6A). These results indicate that both RAB-35 and CED-1 are required for the timely removal of MTM-1 from phagosomal surfaces. Furthermore, *rab-35(b1013)*; *ced-1(n1506)* double mutants display an even longer delay in MTM-1::GFP removal, indicating that *rab-35* and *ced-1* function partially redundantly to regulate MTM-1 removal (Fig 6C). The timing of PtdIns(4,5)P_2_ disappearance and MTM-1 removal from phagosomal surfaces are similar in all backgrounds analyzed (Fig 4D+6C), consistent with the fact that MTM-1 is a PtdIns(4,5)P_2_ effector (25). In addition, it suggests that, in *rab-35* mutants, the persistent presence of PtdIns(4,5)P_2_ on phagosomal surfaces causes MTM-1 to remain on phagosomes as well.

**Fig 6.**
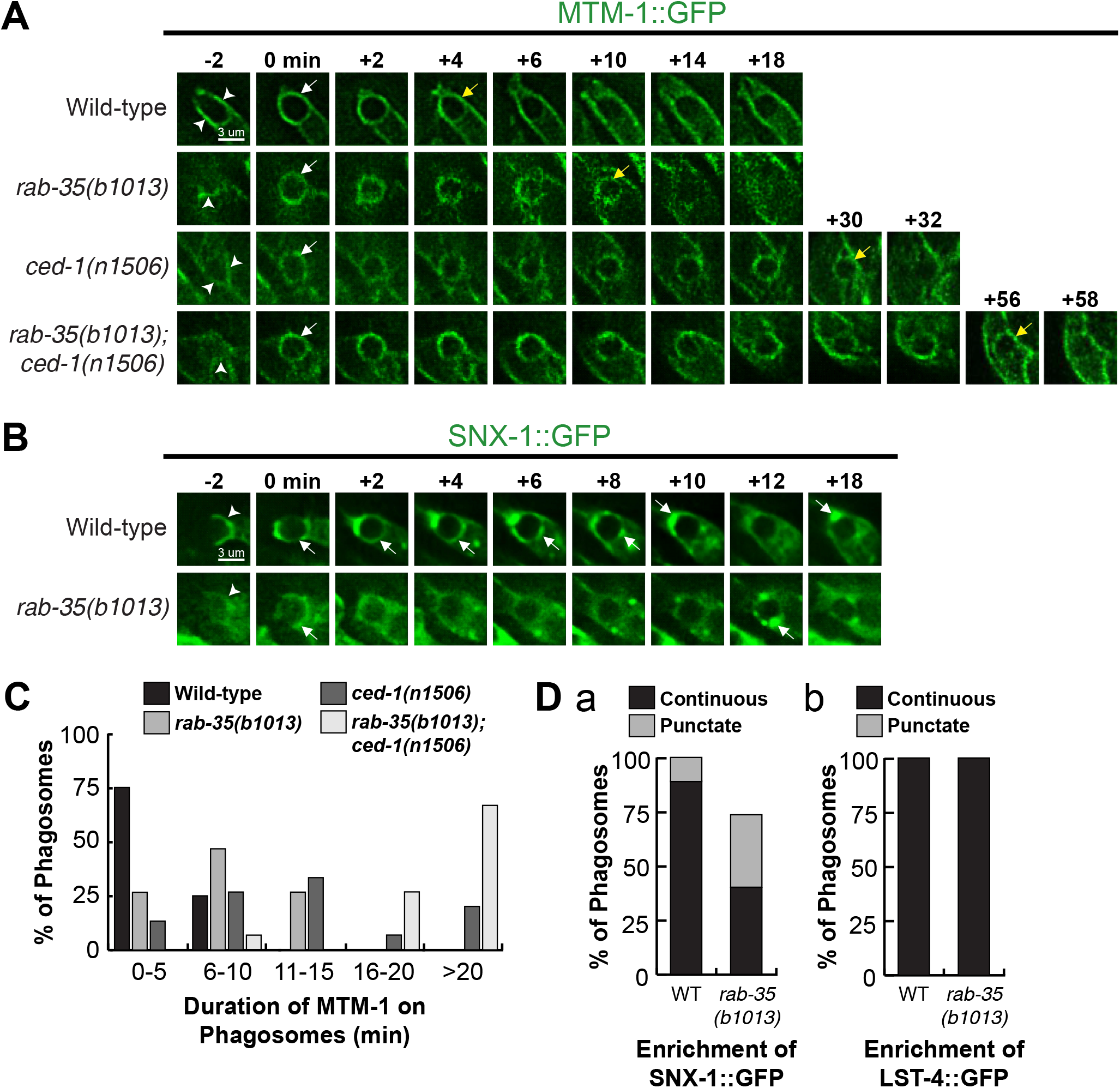
The *rab-35(b1013)* mutation impairs MTM-1 removal from, and SNX-1 recruitment to, phagosomal surfaces. (A) Time-lapse images during and after the formation of a phagosome carrying C3 in embryos of different genotypes expressing P_*ced-1*_ *mtm-1::gfp*. “0 min” indicates the formation of the nascent phagosome. Arrowheads indicate extending pseudopods. White arrows and yellow arrows indicate the time points when MTM-1::GFP first and last appear on the phagosome surface, respectively. (B) Time-lapse images during and after the formation of a phagosome carrying C3 in embryos of different genotypes expressing P_*ced-1*_ *snx-1::gfp*. “0 min” indicates the formation of the nascent phagosome. Arrowheads indicate extending pseudopods. White arrows mark the regions on the phagosomal surface that have an enriched GFP signal. (C) Histogram displaying the range of time that MTM-1 persists on phagosomes in wild-type and *rab-35(b1013)* embryos. Phagosomes bearing cell corpses C1, C2, and C3 were scored. “0 min” indicates the formation of the nascent phagosome. This time interval is defined as that between the formation of a nascent phagosome (“0 min”) and the first time point that MTM-1 is no longer found on the phagosome. For each genotype, at least 15 phagosomes were scored. (D) The efficiency of recruitment of SNX-1::GFP (a) and LST-4::GFP (b) to the surface of phagosomes carrying C1, C2, and C3 was scored in various genotypes. SNX-1::GFP is enriched onto phagosomal surfaces in two different patterns, either distributed onto the entire phagosomal surface evenly (“continuous”) or attached to phagosomal surfaces as puncta (“punctate”) (a), whereas LST-4::GFP is enriched onto phagosomes only in the “continuous” pattern (b). For each genotype, at least 15 phagosomes were scored.

### RAB-35 promotes the recruitment of SNX-1, a PtdIns(3)P effector, to phagosomal surfaces

Sorting nexins SNX-1 and LST-4, two PtdIns(3)P effectors and membrane remodeling factors, are recruited to phagosomal surfaces by PtdIns(3)P (46). SNX-1::GFP and LST-4::GFP were visualized using time lapse microscopy to characterize their localization to phagosomal surfaces. SNX-1::GFP is found on 100% of phagosomes in wild-type embryos; moreover, it forms a continuous ring on the surface in >90% of phagosomes, a pattern suggesting the presence of a large enough number of the SNX-1::GFP molecules to cover the entire surface of a phagosome (Figure 6B+D(a)). In contrast, only 73.3% of phagosomes *rab-35(b1013)* mutants recruit SNX-1::GFP; of these phagosomes, more than half recruit SNX-1::GFP as isolated puncta instead of as a continuous ring (Fig 6B+D(a)). These results suggest that RAB-35 is important for the efficient recruitment of SNX-1::GFP to phagosomes, consistent with the defects in PtdIns(3)P production previously observed in *rab-35* mutants. However, continuous rings of LST-4::GFP were observed on 100% of phagosomes in both *rab-35(b1013)* mutants and wild-type embryos (Fig 6D(b)). Given that our lab has previously shown that the recruitment of LST-4 is mediated in part through DYN-1 (46), this suggests that RAB-35 may also recruit SNX-1 through a more direct mechanism rather than just indirectly through PtdIns(3)P production.

### *rab-35* recruits RAB-5 to phagosomes and acts in the same genetic pathway as *rab-5* during phagosomal maturation

Given that *rab-35(b1013)* mutants are defective in production of PtdIns(3)P, and that the recruitment of RAB-5 and the production of PtdIns(3)P on phagosomal surfaces are co-dependent processes (16), we investigated the functional relationship between RAB-35 and RAB-5. We made a number of observations that indicate that RAB-35 functions upstream of RAB-5 in the regulation of phagosome maturation. Firstly, the enrichment of mRFP::RAB-35 on the surface of nascent phagosomes precedes that of GFP::RAB-5 by approximately 30-60 seconds (Fig 7A). Secondly, inactivation of *rab-5* using RNAi treatment in wild-type embryos results in the presence of extra cell corpses; moreover, the *rab-35* null mutation does not further enhance the Ced phenotype caused by *rab-5*(RNAi) treatment (Fig 7C), suggesting *rab-35* and *rab-5* act in the same genetic pathway. Thirdly, *rab-35(b1013)* mutants exhibit a delay in the recruitment of RAB-5 to the phagosome (Fig 7B+D). This delays resembles that caused by the *ced-1* mutation (Fig 7B+D), although it is not as severe. Additionally, *rab-35(b1013)*; *ced-1(n1506)* double mutants display a stronger delay than either single mutant (Fig 7B+D), suggesting that *rab-35* and *ced-1* function in parallel to recruit RAB-5 to the phagosome.

**Fig 7.**
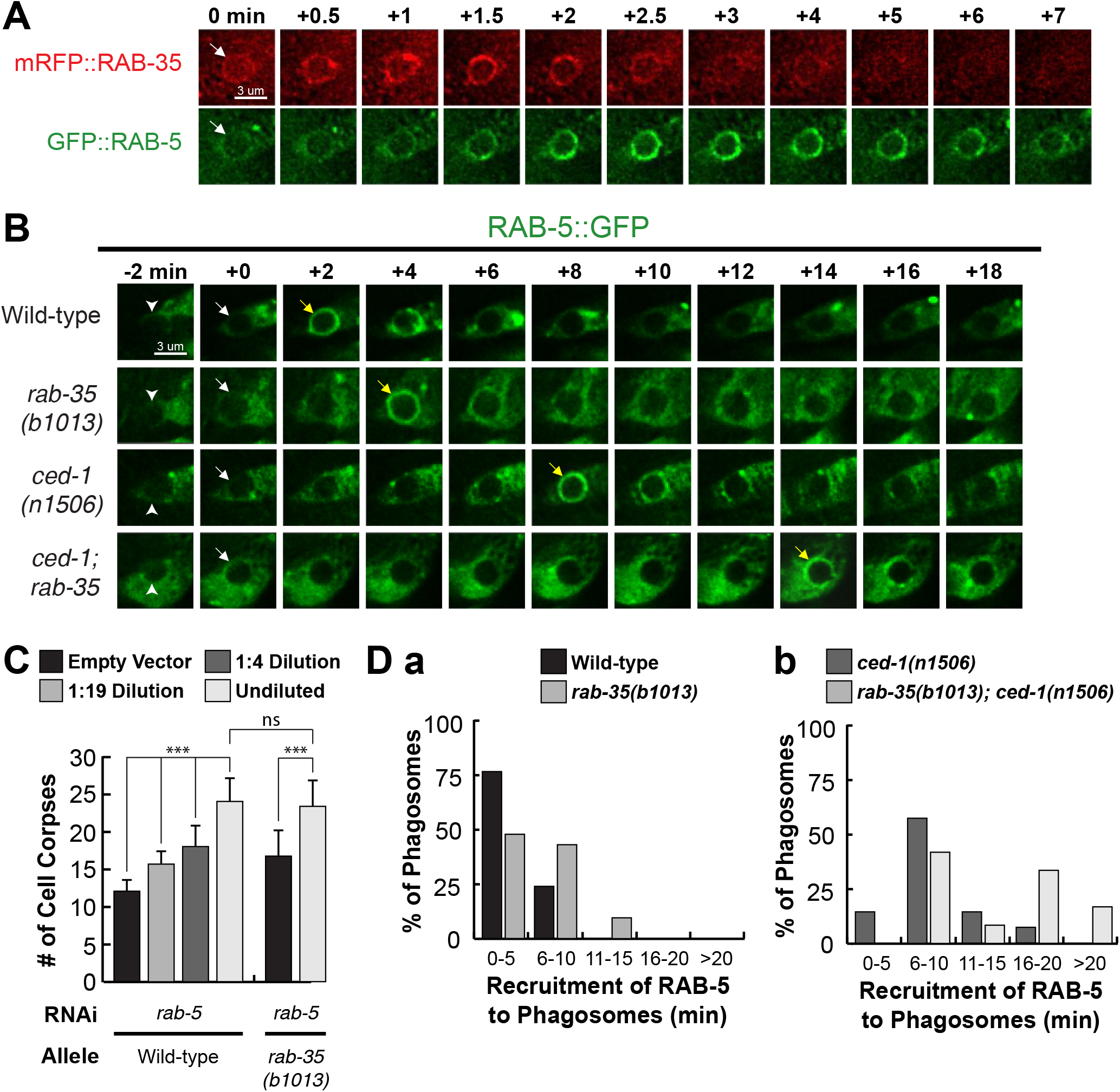
*rab-35* functions upstream of and promotes the phagosomal localization of *rab-5*. (A) Time-lapse images after the formation of a phagosome carrying C3 in a wild-type embryo co-expressing P_*ced-1*_ *mrfp::rab-35* and P_*ced-1*_ *gfp::rab-5*. “0 min” indicates the formation of the nascent phagosome. A whole arrow marks the nascent phagosome. (B) Time-lapse images during and after the formation of a phagosome carrying C3 in a wild-type embryo expressing P_*ced-1*_ *gfp::rab-5*. “0 min” indicates the formation of the nascent phagosome. Arrowheads indicate extending pseudopods. White arrows mark the nascent phagosome, while yellow arrows mark the first time point when RAB-5 localizes to the phagosome. (C) Epistasis analysis was performed between *rab-5* and *rab-35*, using RNAi to inactivate *rab-5*. To analyze the effect of reducing *rab-5* gene dosage, *E. coli* carrying the *rab-5* RNAi construct was diluted by mixing with *E. coli* carrying an empty vector. After RNAi treatment, the numbers of apoptotic cell corpses were scored in the F1 progeny at the 1.5-fold embryonic stage. For each data point, at least 15 embryos were scored. *, 0.001 < p < 0.05; **, 0.00001 < p <0.001; ***, p <0.00001; ns, no significant difference. (D) Histograms displaying the range of time it takes for the appearance of RAB-5 on the surface of phagosomes bearing C1, C2, and C3 in embryos of various genotypes. The histograms exhibit the effect of *rab-35* loss of function in wild-type (a) and *ced-1(n1506)* (b) backgrounds. The time span of RAB-5 appearance is scored as the time interval between the formation of a nascent phagosome (“0 min”) and the first time point that the GFP::RAB-5 signal becomes enriched on the phagosomal surface. For each genotype, at least 15 phagosomes were scored.

### During cell corpse internalization, RAB-35 plays a specific role in the recognition of cell corpses

The initial enrichment of GFP::RAB-35 on extending pseudopods (Fig 2B) suggests that in addition to phagosome maturation, RAB-35 might function in other steps of cell-corpse clearance. To determine whether RAB-35 plays any role in the recognition and/or the engulfment of cell corpses, we took advantage of CED-1ΔC::GFP, a GFP tagged and truncated form of CED-1 that is missing its C-terminal intracellular domain (26). This reporter, when expressed in *ced-1(+)* strains, first clusters to the contact site between the engulfing and dying cell, subsequently spreads to the extending pseudopods, and, when engulfment is complete, finally forms a ring around the nascent phagosome (28). Unlike native CED-1, CED-1ΔC::GFP stays on the surface of a phagosome until it is completely degraded (28). Thus, cell corpses labeled with complete CED-1ΔC::GFP rings must have been previously engulfed, while cell corpses in the middle of being engulfed are labeled with partial GFP^+^ rings that represent phagocytic cups. Cell corpses that are not recognized by an engulfing cell fail to be labeled by a GFP signal (Fig 8A).

**Fig 8.**
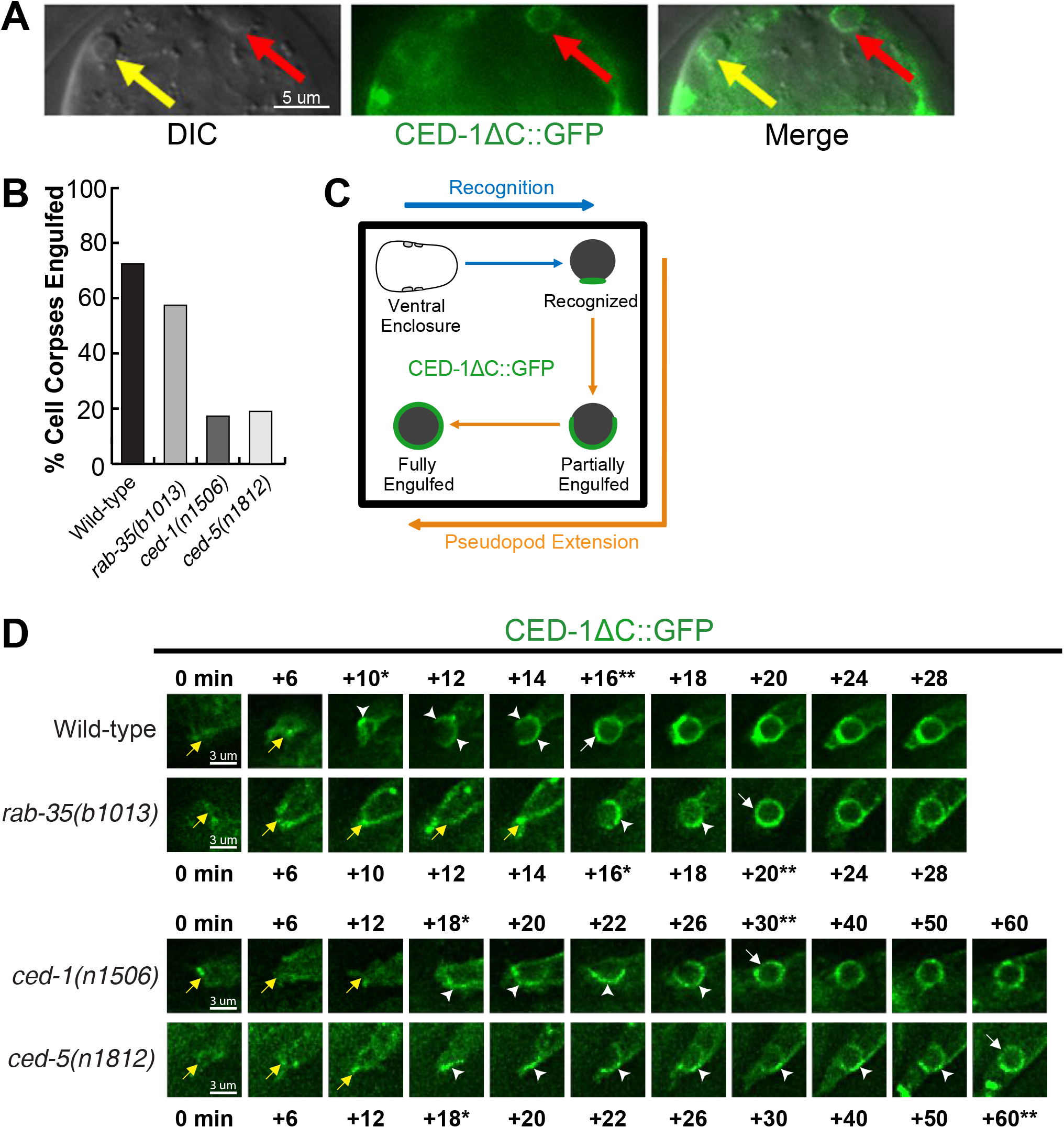
RAB-35, CED-1, and CED-5 function in parallel to engulf apoptotic cell corpses. (A) Images of part of a 1.5-fold stage embryo expressing P_*ced-1*_*ced-1ΔC::gfp*. CED-1ΔC::GFP is utilized to determine whether a cell corpse is engulfed. DIC morphology is used to mark cell corpses. Red arrows indicate an engulfed cell corpse, which is surrounded by a CED-1ΔC::GFP ring. Yellow arrows indicate an unengulfed cell corpse, which lacks CED-1ΔC::GFP on its surface. (B) In 1.5-fold to 2-fold stage embryos of various genotypes, the fraction of cell corpses that had been engulfed was measured using the CED-1ΔC reporter. For each genotype, at least 15 embryos were scored. (C) Diagram outlining the assays used to determine the moments of cell corpse recognition C::GFP reporter. The moment of recognition is defined as the first time point GFP is seen enriched in a region in contact between the engulfing and dying cell, with the moment of ventral enclosure used as a reference point (“0 min”). The period of pseudopod extension is defined as the time span between the moment of recognition and the moment when the nascent phagosome forms. (D) Time-lapse images before and after the formation of a phagosome carrying C3 in embryos of different genotypes expressing P_*ced-1*_*ced-1ΔC::gfp* “0 min” indicates the initiation of ventral enclosure. Arrowheads indicate extending pseudopods. A white arrow marks the nascent phagosome. Yellow arrows label the extending ventral hypodermal cell ABpraapppp. For each genetic background, a single asterisk marks the time point when recognition is first observed, while two asterisks marks the time point when the nascent phagosome is formed.

We first analyzed all cell corpses in mid-stage (1.5-fold stage) embryos. In *rab-35(b1013)* embryos, a significantly lower percentage of engulfed cell corpses were observed compared to wild-type embryos (Fig 8B), suggesting that the *rab-35* mutation causes defects in the internalization of cell corpses. Such a failure in cell corpse internalization may result from defects in the recognition or the actual engulfment of the cell corpse. To distinguish which of these is the case, we monitored the generation and extension of pseudopods around dying cells C1, C2, and C3 using the CED-1ΔC::GFP reporter. In this assay, a delay in the clustering of GFP signal on the site of contact between a cell corpse and its engulfing cell indicates a delay in the recognition of the cell corpse. To further discern the precise moment that pseudopod extension initiates, we took advantage of the temporal consistency of *C. elegans* development between embryos, marking the moment when the two ventral hypodermal cells ABplaapppp and ABpraapppp begin to extend towards the ventral midline as the “0” time point (Fig 8C). Because P_*ced-1*_ *ced-1ΔC::gfp* is expressed in embryonic hypodermal cells and localizes to the plasma membrane, the GFP signal allows us to accurately record this moment (Fig 8D).

We observed that the recognition of cell corpses was delayed in *rab-35(b1013)* mutant embryos (Fig S5). In wild-type embryos, 40% of the cell corpses are recognized within the first 10 minutes of ventral enclosure, yet in *rab-35(b1013)* mutants, only 6.7% cell corpses are recognized within that same time period (Fig S5A(a)). Additionally, in *rab-35(b1013)* mutants, 20% of the cell corpses are recognized between 21-30 minutes after the start of ventral enclosure, whereas only 6.7% of cell corpses take that long to be recognized in wild-type embryos (Fig S5A(a)). Conversely, *rab-35* loss of function causes no observable delays in pseudopod extension or phagosome sealing once the cell corpse is recognized (Fig S5B). Together, these results indicate that during the cell-corpse internalization process, RAB-35 specifically regulates the recognition of cell corpses.

### RAB-35 acts in a pathway separate from the CED-1 or CED-5 pathways to promote the recognition of cell corpses

We found that null mutants of *ced-1* and *ced-5*, two engulfment genes that each represents one of the two parallel pathways for engulfment (8–47,48), display significantly greater delays in cell corpse recognition relative to *rab-35(b1013)* null mutants (S5A(a-c)), indicating that both CED-1 and CED-5 are essential for the timely recognition of apoptotic cells. The *ced-1; ced-5* double null mutant embryos suffer greater recognition delay than each single mutant strain (Fig S5A(b-d)), supporting this conclusion. We further observed that in double mutant combinations, *rab-35(b1013)* mutants enhanced the recognition delay of both *ced-1(n1506)* and *ced-5(n1812)* mutants (Fig S5A(b-c)), suggesting that *rab-35* acts in a previously unknown pathway separate from the *ced-1* or *ced-5* pathways in the context of cell corpse recognition. The *ced-1; rab-35; ced-5* triple null mutants suffer from a stronger recognition delay than any of the respective double mutants, with 63.2% of cell corpses being recognized more than 40 minutes after ventral enclosure started (Fig S5A(d)), further supporting our hypothesis.

### RAB-35 represents a third genetic pathway that facilitates the clearance of cell corpses in parallel to the CED-1/6/7 and CED-2/5/10/12 pathways

We have identified the functions of RAB-35 in two distinct cell-corpse clearance events: (I) the recognition and (II) the degradation of cell corpses. In both of these events, RAB-35 appears to function independently of both CED-1 and CED-5. To further determine whether *rab-35* represents a third pathway in addition to the *ced-1* and *ced-5* pathways to regulate apoptotic cell clearance, we performed a thorough epistasis analysis between *rab-35* and the two canonical pathways, the *ced-1*/*ced-6*/*ced-7* pathway and the *ced-2*/*ced-5*/*ced-10*/*ced-12* pathway (48). This epistasis analysis was performed by quantifying the number of cell corpses in double mutant combinations between the null allele *rab-35*(*b1013)* and null alleles of representative genes in each of the two canonical engulfment pathways. Remarkably, *rab-35* was found to be parallel to multiple components of both the *ced-1*/*ced-6*/*ced-7* pathway (*ced-1* and *ced-6*) and the *ced-2*/*ced-5*/*ced-10*/*ced-12* pathway (*ced-5*, *ced-10*, and *ced-12*); *rab-35* loss of function tremendously enhances the Ced phenotype of each of these mutants in 1.5-fold embryos, 4-fold embryos, and the 48-hour post-L4 adult gonad (Fig 9A-B). In *rab-35(b1013); ced-1(n1506)*; *ced-5(n1812)* triple mutants, 4-fold embryos contain nearly as many apoptotic cell corpses as 1.5-fold embryos, suggesting that cell corpses produced throughout embryogenesis persist until hatching; this behavior is indicative of a near complete block of apoptotic cell clearance (Fig 9A-B).

**Fig 9.**
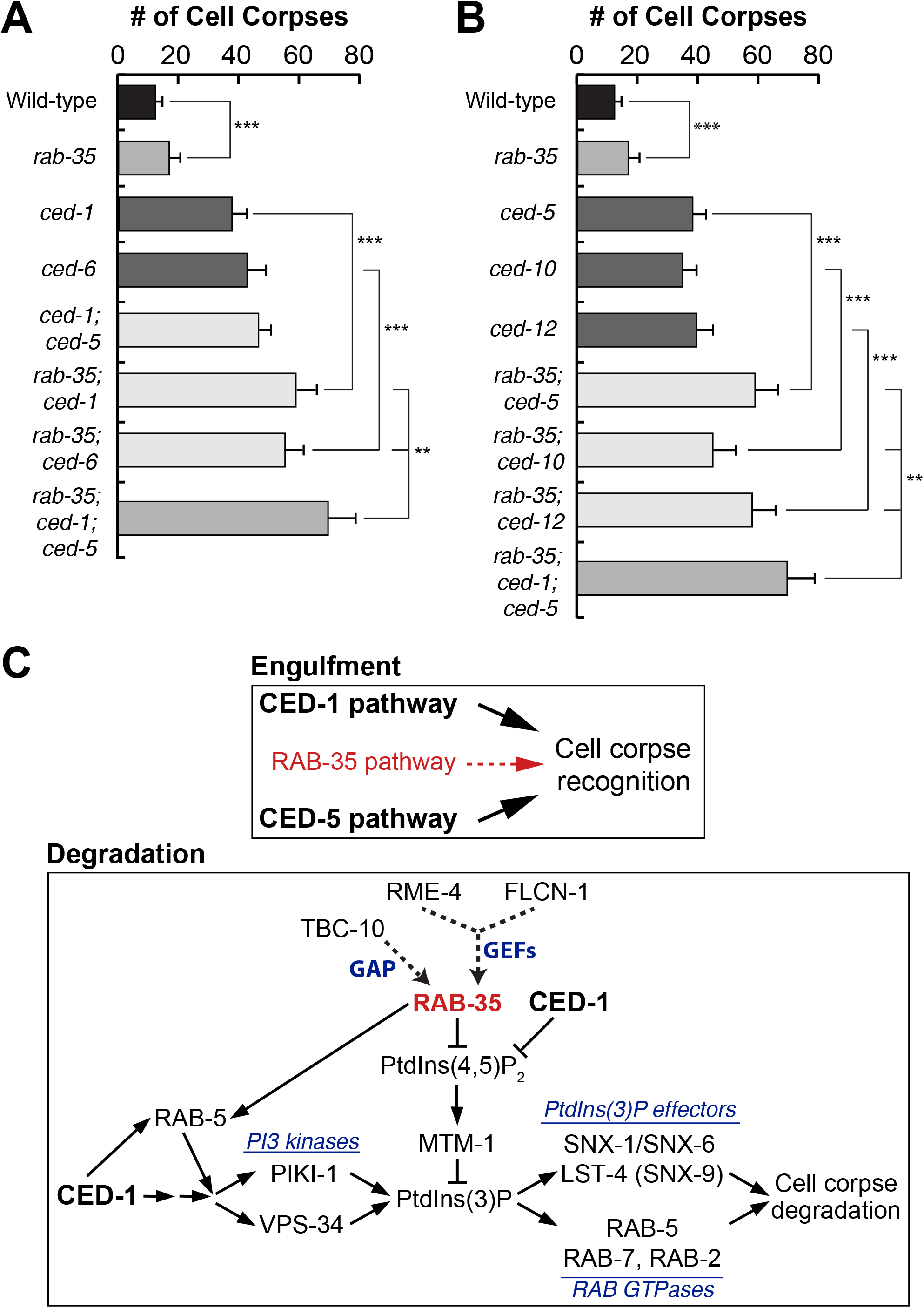
*rab-35* represents a third engulfment pathway independent of both the *ced-1/-6/-7* and *ced-2/-5/-10/-12* pathways. The average numbers of cell corpses in 1.5-fold stage embryos of various genotypes are presented as bars in the bar graphs. Error bars indicate sd. The student *t*-test was used for data analysis: *, 0.001 < p < 0.05; **, 0.00001 < p <0.001; ***, p <0.00001; ns, no significant difference. For each data point, at least 15 animals were scored. (A) Results of epistasis analysis performed between *rab-35* and components of the *ced-1/-6/-7* pathway. Null alleles [*rab-35(b1013)*, *ced-1(n1506)*, and *ced-6(n2095)*] were used. (B) Results of epistasis analysis performed between *rab-35* and components of the *ced-2/-5/-10/-12* pathway. Null alleles [*ced-5(n1812)* and *ced-12(n3261)*] were used; however, null alleles of *ced-10* (*C. elegans* ortholog of mammalian Rac1) are embryonic lethal, so a severe loss-of-function allele (*n1993*) was used instead. (C) Diagram illustrating the role of RAB-35 in the engulfment and degradation of cell corpses in *C. elegans*. Please see Discussion for more details. RAB-35 functions in parallel with the *ced-1/-6/-7/dyn-1* and *ced-2*/*-5/-10*/*-12* pathways in the recognition of cell corpses, while RAB-35 and CED-1 function in parallel during phagosome maturation. RAB-35 helps to recruit RAB-5, which promotes the production of PtdIns(3)P. Furthermore, RAB-35 stimulates the turnover of both PtdIns(4,5)P_2_ and its effector MTM-1, a PI-3 phosphatase. In turn, PtdIns(3)P promotes the recruitment of the sorting nexins SNX-1, SNX-6, and LST-4 as well as the Rab GTPases RAB-2, RAB-5, and RAB-7, allowing for the progression of phagosome maturation and cell corpse degradation. Among the three sorting nexins, *rab-35* mutants are defective in the recruitment of SNX-1 but not LST-4/SNX-9 to phagosomes, suggesting that the effect might be too weak to measure and/or that other factors may be involved in their recruitment.

## Discussion

The study of apoptotic cell clearance is still relatively nascent, and a number of important components – such as *C. elegans* RAB-35 – are in the process of being discovered. Rab35 is a multifunctional GTPase that plays important roles in a wide variety of biological processes. Mammalian Rab35 has been implicated in events including, but not limited to: exocytosis (49,50), endocytic recycling (51–54), cytokinesis (55,56), cytoskeleton rearrangement (56–61), and autophagy (62). Recently, Rab35 was found to be an oncogene that promotes proliferation by activating the PI3K/AKT signaling pathway (63). Like its mammalian homolog, *C. elegans* RAB-35/RME-5, as well as its GEF RME-4, were found to act in endocytic recycling and yolk uptake in the developing oocytes (41).

Mammalian and *Drosophila* Rab35 have been implicated in phagocytosis and phagosome maturation, two processes that are closely linked with apoptotic cell clearance. Inactivating Rab35 reduces the internalization efficiency of macrophages against erythrocytes, zymosan particles, and microbes (57,64–66). In addition, overexpression of dominant negative Rab35(S22N) inhibits the maturation of phagosomes carrying pathogenic bacteria (67). However, besides finding that Rab35 facilitates phagocytic cup formation through the ARF6 GTPase, which in turn regulates the actin cytoskeleton (64,65), not much else is known about the molecular mechanisms employed by Rab35 to support phagosome formation and maturation. In addition, whether Rab35 plays any role in the clearance of apoptotic cells was not known.

Our work in *C. elegans* has discovered that RAB-35 regulates multiple cell corpse clearance events (Fig 9C). We have uncovered a novel role in the recognition of cell corpses by RAB-35, a process that enables engulfment and the formation of a nascent phagosome. RAB-35 then helps to initiate the maturation of this nascent phagosome through novel molecular mechanisms that promote the PtdIns(4,5)P_2_ to PtdIns(3)P switch on phagosomes and aid in the recruitment of RAB-5 to the phagosomal surface. Furthermore, RAB-35 leads a genetic pathway in parallel to the two known pathways for apoptotic cell clearance, establishing mechanisms that promote robustness and ensure efficiency throughout clearance.

### RAB-35 function depends on its cycling between GDP- and GTP-bound forms, a process facilitated by the GAP TBC-10 and the GEFs FLCN-1 and RME-4

For many Rab small GTPases, such as Rab7, the GTP- and GDP-bound forms are their active and inactive forms, respectively (18,68,69). Other small GTPases, such as RAB-5, need to cycle between the GTP- and GDP-bound forms in order to function (70). Our characterization of the presumed GTP- and GDP-locked mutant forms of RAB-35 has revealed that the specific function of RAB-35 in apoptotic cell clearance depends on the cycling between its GTP- and GDP-bound forms, resembling the dynamics observed in RAB-5.

Among three *C. elegans* homologs of mammalian proteins known to act as Rab35 GAPs (71), we have identified TBC-10 as the GAP for RAB-35 in the context of apoptotic cell clearance. In *tbc-10* deletion mutants, where RAB-35 supposedly is locked in the GTP-bound state, GFP::RAB-35 persists on the surfaces of phagosomes, yet apoptotic cell clearance is defective in a manner akin to *rab-35* null mutants. These observations support the hypothesis that RAB-35 must cycle between its GTP- and GDP-bound forms to properly function, although the reasons behind this phenomena are yet unclear.

Although the *in vitro* GEF activity of folliculin towards mammalian Rab35 has been detected (72), and folliculin was reported to activate Rab35 in order to mediate EGF receptor recycling in a cancer cell line (73), the functional relationship between folliculin and Rab35 under physiological conditions in an animal context has not been reported. We found that *flcn-1*, the *C. elegans* homolog of folliculin, acts in the same genetic pathway as *rab-35* does to promote cell-corpse clearance, suggesting that FLCN-1 is a putative GEF for RAB-35 during cell-corpse clearance. This is the first time that folliculin has been implicated as a GEF for RAB-35 during animal development.

While *flcn-1* and *rab-35* act in the same genetic pathway, the Ced phenotype displayed by the *flcn-1* null mutants is slightly weaker than that of the *rab-35* null mutants. We observed that *C. elegans* RME-4, a homolog of the connecdenns DENND1A-C and a GEF for RAB-35 in yolk receptor recycling (41), activates RAB-35 alongside FLCN-1 during apoptotic cell clearance. However, the *rme-4* null mutation does not cause a significant defect in clearance by itself, suggesting that FLCN-1 acts as the predominant GEF for RAB-35 in the context of apoptotic cell clearance, while RME-4 only plays a minor role. Our results are consistent with the observation that, as a multifunctional GTPase, RAB-35 is regulated by different GEFs in each cellular event that it is involved (44).

### RAB-35 modulates the initiation of phagosome maturation by regulating phosphatidylinositol dynamics and RAB-5 recruitment

Phosphorylated forms of phosphatidylinositol species are second messengers that play essential roles leading the formation and degradation of phagosomes (24). During apoptotic cell clearance in *C. elegans*, a process known as the PtdIns(4,5)P_2_ to PtdIns(3)P switch occurs immediately after the sealing of pseudopods and the formation of nascent phagosomes. During this shift, PtdIns(4,5)P_2_ – which had been enriched on extending pseudopods – rapidly disappears from phagosomal surfaces, while PtdIns(3)P – essential for the initiation of phagosome maturation – subsequently appears on phagosomal surfaces at a high level and oscillates in a biphasic pattern (16,25).

We have observed that once engulfment starts, GFP::RAB-35 becomes enriched on the surface of extending pseudopods. The pseudopod localization pattern overlaps with that of PtdIns(4,5)P_2_ and can be explained by its membrane-anchoring prenylation motif typical of Rab GTPases (74) as well as by an evolutionarily conserved polybasic region that has a high affinity for negatively charged phosphatidylinositol species such as PtdIns(4,5)P_2_ (75–77). Immediately after engulfment, RAB-35 is further transiently enriched on the surface of the nascent phagosome. On nascent phagosomes, the initiation of this RAB-35 enrichment coincides perfectly with both the turnover of PtdIns(4,5)P_2_ and the appearance of PtdIns(3)P. This unique pattern is consistent with a role of RAB-35 in the switch of phagosomal phosphatidylinositol species from PtdIns(4,5)P_2_ to PtdIns(3)P.

Furthermore, we found that *rab-35* mutants suffer significant delays in both the disappearance of PtdIns(4,5)P_2_ and the appearance of the first wave of PtdIns(3)P on phagosomal membranes. Interestingly, we found no defects in the recruitment of the PI3-kinase PIKI-1. However, we discovered that the PI3-phosphatase MTM-1, a PtdIns(4,5)P_2_ effector that dephosphorylates PtdIns(3)P and in this way counteracts the function of PI3-kinases, persists on the surface of nascent phagosomes much longer in *rab-35* mutants. Together, the above evidence indicates that RAB-35 promotes the turnover of PtdIns(4,5)P_2_ on phagosomal membranes, which in turn removes MTM-1 and consequently suppresses PI3-phosphatase activity. We have also found that RAB-35 contributes to RAB-5 recruitment on phagosomal surfaces. As RAB-5 promotes the production of PtdIns(3)P on phagosomal surfaces (16), we propose that RAB-35 facilitates the robust production of PtdIns(3)P on phagosomal surfaces through two separate activities, the removal of PtdIns(4,5)P_2_ and the recruitment of RAB-5 (Fig 9C).

This delay of the robust production of PtdIns(3)P observed in *rab-35* mutants is associated with numerous defects in phagosome maturation: (I) The degradation of cell corpses as a whole is delayed; (II) The recruitment of early endosomes, an intracellular organelle that is incorporated into phagosomes and is essential for phagosome maturation (13), is also delayed; and (III) SNX-1, a sorting nexin and a PtdIns(3)P effector known to promote phagosome maturation (46), is recruited to the phagosomal surface less efficiently. Considering that SNX-1 is necessary for the recruitment of early endosomes to phagosomes (46), we propose that this defect in SNX-1 recruitment observed in *rab-35* mutants causes a defect in the recruitment of early endosomes. Our findings thus uncover a novel molecular mechanism employed by RAB-35 for cell-corpse degradation and delineates a novel pathway led by RAB-35 that is responsible for regulating the PtdIns(4,5)P_2_ to PtdIns(3)P switch, an event essential for the initiation of phagosome maturation.

What we report suggests that RAB-35 promotes phagosome maturation primarily through the removal of PtdIns(4,5)P_2_ on phagosomal surfaces. In addition, inactivation of mammalian Rab35 results in the accumulation of PtdIns(4,5)P_2_ on intracellular vacuoles and other structures (55,58), suggesting that the timely turnover of PtdIns(4,5)P_2_ is a conserved activity of Rab35. How, then, does the localization and activation of RAB-35 on the surface of a nascent phagosome cause the disappearance of PtdIns(4,5)P_2_? The level of PtdIns(4,5)P_2_ is known to be determined by two antagonizing activities, the phosphorylation of PtdIns(4)P or PI(5)P by PtdIns kinases and the dephosphorylation of PtdIns(4,5)P_2_ by PtdIns phosphatases (78). OCRL, a 5-phosphatase that dephosphorylates PtdIns(4,5)P_2_, physically interacts with the GTP-bound form of Rab35 and was reported to function as a Rab35 effector that regulates PtdIns(4,5)P_2_ turnover during cytokinesis (58). OCRL-1, its *C. elegans* homolog, was reported to play an active role in modulating PtdIns(4,5)P_2_ turnover on the surfaces of pseudopods and phagosomes, and its inactivation results in a strong Ced phenotype indicative of a cell corpse clearance defect (25). Thus, we propose that OCRL-1 is a promising candidate linking RAB-35 with PtdIns(4,5)P_2_ levels on the phagosomal surface; however, whether RAB-35 regulates the activity and/or subcellular localization of OCRL-1 has yet to be investigated.

### RAB-35 acts as a robustness factor and defines a novel third cell corpse-clearance pathway

The phagocytic receptor CED-1 and its adaptor CED-6 lead a phagosome maturation pathway in addition to their known role in the recognition of cell corpses (13). This pathway promotes PtdIns(3)P production on – and Rab GTPase recruitment to – phagosomal surfaces (13). We observed that all of the defects in phagosome maturation displayed by *rab-35* null mutants, including the persistence of PtdIns(4,5)P_2_ and its effector MTM-1 on nascent phagosomes, the delay in phagosomal PtdIns(3)P production, the various defects in the recruitment of SNX-1 and RAB-5, and the delay of the incorporation of early endosomes to phagosomes, are also displayed by *ced-1* null mutants (8,20,46) (Figs 4–6). However, all of these defects are much more severe in *ced-1* mutants; for instance, phagosomal PtdIns(3)P production, the incorporation of early phagosomes, and phagosome degradation are frequently blocked (8,18), whereas in *rab-35* mutants they are merely delayed. To determine whether RAB-35 functions as a component of the CED-1 pathway during phagosome maturation, we analyzed *rab-35; ced-1* double mutants. To our surprise, the double mutants display enhanced defects relative to their single mutants in all of the assays mentioned above, indicating that RAB-35 initiates phagosome maturation in a manner independent of the CED-1 pathway. Given that CED-1 controls other events such as the incorporation of lysosomes to phagosomes in addition to those events regulated by RAB-35 (18), our observations suggest that this novel RAB-35 pathway acts redundantly to the CED-1 pathway in some, but not necessarily all, functions of phagosome maturation.

Two parallel pathways, the *ced-1/-6/-7* and *ced-2/-5/-10/-12* pathways, are known to control the recognition and engulfment of cell corpses (18,48,79). The *rab-35(b1013)* null mutation also causes a delay during this process, but this defect is not as severe as that observed with either the *ced-1* or *ced-5* null mutations. While *rab-35* mutants exhibit no defects in either pseudopod extension or phagosome sealing, *rab-35* loss of function enhances the defect in cell corpse recognition observed in *ced-1* mutants, suggesting that RAB-35 and CED-1 function in parallel on engulfing cells to recognize apoptotic cells. Furthermore, when we perform epistasis analysis to measure the overall clearance defect by counting the number of persistent cell corpses in double and triple mutant embryos, we found that *rab-35* defines a novel third pathway by acting in parallel to both the *ced-1/-6/-7* and *ced-2/-5/-10/-12* pathways. Currently, the identity of the phagocytic receptor(s) acting in this third pathway remains unknown, although we strongly suspect transmembrane receptors known as integrins because they have been previously implicated in apoptotic cell clearance in both mammals and *C. elegans* (37,38,80,81). Further investigation is needed to determine whether any integrins act in the *rab-35* pathway.

Given that all phenotypes observed in *rab-35* single mutants are relatively modest, what is the purpose of this novel *rab-35* pathway during apoptotic cell clearance? We propose that RAB-35 acts as a robustness factor that provides a “buffer” to maintain the stability and effectiveness of cell corpse clearance. Robustness factors are important in maintaining system stability when animals encounter genetic or environmental changes. Indeed, after *rab-35* mutant embryos are subject to heat treatment, the Ced phenotype is further enhanced (Fig S6), indicating that RAB-35 helps to keep the mechanisms behind apoptotic cell clearance stable when the system is stressed. When both the CED-1 and CED-5 pathways are intact, missing RAB-35 activity only causes modest defects, much weaker than missing either of the two canonical pathways; however, when one or both of the two canonical pathways is inactivated or when under stress, the RAB-35 pathway provides the necessary activity to support cell corpse clearance activity. Considering that diseases can be thought of a subversion of the “robust yet fragile” nature of optimized and complex biological systems (82–84), we postulate that RAB-35 plays a critical role in health. This effect is likely enhanced in aging individuals that experience an increased incidence of autoimmunity and cancer (85,86), which are associated with defects in apoptotic cell clearance and RAB-35 function (3,63,71,87). Further exploring this physiological role for RAB-35 will help broaden our view of the function of Rab GTPases in both development and diseases.

## Materials and Methods

### Mutations, strains, and transgenic arrays

*C. elegans* strains were grown at 20°C as previously described (88). The N2 Bristol strain was used as the reference wild-type strain. Mutations and integrated arrays are described by Riddle et al. (1997) and the Worm Base (http://www.wormbase.org), except when noted otherwise: LGI, *ced-1(n1506)*, *ced-12(n3261)*, *rab-2/unc-108(n3263), rab-10(ok1494), unc-75(e950)*; LGII, *flcn-1(ok975), rab-7(ok511)*; LGIII, *ced-6(n2095), rab-35(b1013, tm2058), tbc-10(tm2790)*; LGIV, *ced-5(n1812), ced-10(n1993)*; LGV, *unc-76(e911)*; LGX, *rme-4(ns410), tbc-13(ok1812)*. All *ok* alleles were generated by the *C. elegans* Gene Knockout Consortium and distributed by *Caenorhabditis* Genetics Center (CGC). All *tm* alleles were generated and provided by the National Bioresource Project of Japan. Transgenic lines were generated by microinjection as previously described (89). Plasmids were injected alongside the coinjection marker pUNC76 [*unc-76(+)*] into *unc-76(e911)* mutant adult hermaphrodites as previously described (90), with non-Unc animals being identified as transgenic animals.

### Plasmid construction

The cDNAs for *rab-11.1, −18, −19, −30. −33, −35, glo-1, and nuc-1* were amplified from a mixed-stage *C. elegans* cDNA library (Z. Zhou and H.R. Horvitz, unpublished data) using polymerase chain reaction (PCR). The cDNAs for *rab-11.1, −18, −19, −30. −33, −35, glo-1* were cloned into RNAi-by-feeding vector L4440 to generate RNAi constructs. P_*ced-1*_ *gfp::rab-35* was constructed by cloning the *rab-35* cDNAs into the XmaI and KpnI sites of pZZ956 (P_*ced-1*_ *gfp*). P_*ced-1*_ *mrfp::rab-35* was constructed by replacing the *gfp* cDNA in P_*ced-1*_ *gfp::rab-35* with *mrfp* cDNA. The (S24N) and (Q69L) mutations were introduced into P_*ced-1*_ *gfp::rab-35* using the QuickChange Site-directed Mutagenesis Kit (Stratagene, La Jolla, CA) to generate P _*ced-1*_ *gfp::rab-35(S24N)* and P_*ced-1*_ *gfp::rab-35(Q69L)*, respectively. Using the same kit, the S33N mutation was introduced into P_*hsp-16/2*_ *rab-5* and P_*hsp-16/41*_ *rab-5*. P_*hsp-16/2*_ *gfp::rab-5(S33N)* and P_*hsp-16/41*_ *gfp::rab-5(S33N)* were produced by inserting the gfp cDNA into P_*hsp-16/2*_ *rab-5(S33N)* and P_*hsp-16/41*_ *rab-5(S33N)*, respectively. The *nuc-1* cDNA was inserted into the BamHI and XmaI sites of pZZ829 (P_*ced-1*_ *gfp*) to generate P_*ced-1*_ *nuc-1::gfp*. P_*ced-1*_ *nuc-1::gfp* was generated by replacing the *gfp* cDNA with the *mrfp* cDNA. All plasmids contain an *unc-54* 3’ UTR.

### RNA interference (RNAi)

RNAi screen of the candidate *rab* genes was performed using the feeding protocol as previously described (91). The RNAi feeding constructs for *rab-11.1*, *-18*, *-19*, *-30*, *-33*, *-35*, and *glo-1* were produced by our lab, while the remaining constructs came from a *C. elegans* RNAi library (92,93). Mid-L4 stage hermaphrodites were placed on plates seeded with *E. coli* containing the RNAi feeding construct. After 48 hrs, the numbers of germ cell corpses per gonad arm were scored using a DIC microscope.

RNAi of *rab-5*, *ina-1*, *pat-2* was performed using the same feeding protocol as above using constructs from the same library except that, 24 hrs after L4-stage hermaphrodites were placed on RNAi feeding plates, these adults were transferred to a second RNAi plate. After an additional 24 hours, the numbers of cell corpses in 1.5-fold and late 4-fold stage embryos were scored using a DIC microscope.

### Nomarski DIC microscopy

DIC microscopy was performed using an Axionplan 2 compound microscope (Carl Zeiss, Thornwood, NY) equipped with Nomarski DIC optics, a digital camera (AxioCam MRm; Carl Zeiss), and imaging software (AxioVision; Carl Zeiss). Previously established protocols were used to score cell corpses under DIC microscopy (8,43). Somatic embryonic cell corpses were scored in the head region of embryos at various developmental stages (comma, 1.5-fold, 2-fold, late 4-fold, and early L1). Germ cell corpses were scored in one of the two gonadal arms of adult hermaphrodites 24 or 48 hrs after the mid-L4 stage. Yolk analysis was performed by characterizing the amount of yolk found in the pseudocoelom near the gonads of adult hermaphrodites 24 or 48 hrs after the L4 stage.

### Fluorescence microscopy and quantification of cell corpse clearance events

An Olympus IX70-Applied Precision DeltaVision microscope equipped with a DIC imaging apparatus and a Photometris Coolsnap 2 digital camera was used to capture fluorescence and DIC images, while Applied Precision SoftWoRxV software was utilized for image deconvolution and processing (43). To quantify the number of engulfed cell corpses in 1.5-fold to 2-fold stage embryos expressing CED-1ΔC::GFP, both DIC and GFP images of 40 serial z-sections at a 0.5-μm were recorded for each embryo. Engulfed cell corpses were those labeled with a full GFP^+^ circle. Unengulfed cell corpses were those that display the refractive appearance under DIC optics yet were either labeled with a partial GFP^+^ circle or not labeled at all.

The dynamics of various GFP, mRFP, and mCherry reporters during the engulfment and degradation of cell corpses C1, C2, and C3 were examined using an established time-lapse recording protocol (18,43). Ventral surfaces of embryos were initially monitored 300-320 minutes post-first cleavage. Recordings typically lasted 60-180 minutes, with an interval of 30 secs to 2 mins. At each time point, 10-16 serial z-sections at a 0.5-μm interval were recorded. Signs such as embryo elongation and embryo turning prior to comma stage were closely monitored under DIC to ensure that the embryo being recorded was developing normally. The moment of cell corpse recognition is the time when CED-1ΔC:GFP first clusters to the region where an engulfing cell contacts a cell corpse, measured relative to the moment ventral enclosure begins; the initiation of ventral enclosure is defined as the time point when hypodermal cells ABplaapppp and ABpraapppp begin to extend across the ventral surface. The time period of pseudopod extension is the time interval between when budding pseudopods labeled with CED-1ΔC::GFP are first observed and when the two pseudopods join and seal to form a nascent phagosome. The life span of a phagosome is defined as the time interval between when pseudopods seal to form the nascent phagosome and when the phagosome shrinks to one-third of its original radius.

## Acknowledgements

We thank K. Venken for reagents for Gibson cloning. We thank D. Reiner, S. Sazer, K. Venken, M. Wang, and T. Wensel for advice and helpful comments. We thank the Caenorhabditis Genetics Center (CGC) and the National BioResource Project in Japan (Shohei Mitani) for providing mutant strains.

## Author Contributions

The author(s) have made the following declarations about their contributions: Conceived and designed the experiments: RH ZZ. Performed the experiments: RH YW ZZ. Analyzed the data: RH YW ZZ. Wrote the paper: RH ZZ.

## Supplemental Figure Legends

**Fig S1. Loss of function of *rab-35* causes the appearance of excess yolk in the pseudocoelom.**

(A) Differential interference contrast (DIC) microscopy images of adult gonads of (a) wild-type and (b) *rab-35(b1013)* mutants. White arrows mark pools of yolk. *rab-35(b1013)* mutants contain excess yolk in the pseudocoelom compared to wild-type. (B) Wild-type and *rab-35(b1013)* mutant worms were visualized as adults using DIC microscopy, characterized based on the yolk coverage, and separated into four distinct groups. (C) Wild-type and *rab-35(b1013)* were scored as 24-hour and 48-hour post-L4 adults based on yolk coverage.

**Fig S2. TBC-10, RME-4, and FLCN-1 are orthologs of mammalian TBC1D10A, connecdenn 1/2/3, and folliculin, respectively.**

Homology between TBC-10 and its human ortholog, TBC1D10A. TBC-10 and TBC1D10A share 29.6% identity and 42.3% similarity overall, and share 61.6% identity and 73.5% similarity within the highly conserved TBC (Tre-2/Bub2/Cdc16) GAP domain. The TBC domain is highlighted in yellow. All alignments were performed using EMBOSS Needle. Asterisks (*) indicate identical amino acids, colons (:) indicate similar substitutions, periods (.) indicate non-similar substitutions, and dashes (−) indicate areas where no alignment was possible. The residues absent in *tbc-10(tm2790)* mutants are highlighted in red, while residues absent in *tbc-10(tm2907)* are highlighted in blue. Homology between the first 500 residues of RME-4 and its human orthologs, DENND1A/connecdenn 1, DENND1B/connecdenn 2, and DENND1C/connecdenn 3 [only DENND1A is shown]. RME-4 shares 22.5% identity and 34.9% similarity overall with DENND1A; 23.6% identity and 37.4% with DENND1B; and 26.4% identity and 40.2% similarity with DENND1C. Within the more highly conserved DENN (differentially expressed in normal and neoplastic tissue) GEF domain, these values increase to 41.0% identity/67.6% similarity; 40.3% identity/66.9% similarity; and 41.7%/65.5%, respectively. The uDENN (upstream of DENN) domain is highlighted in blue, the DENN domain is highlighted in yellow, and the dDENN (downstream of DENN) domain is highlighted in green. The residues absent in *rme-4(tm1865)* mutants are highlighted in red. (C) Homology between FLCN-1 and its human ortholog folliculin. FLCN-1 and folliculin have non-canonical DENN domains, and unlike their counterparts found within RME-1 and DENND1A/B/C, they are not specifically conserved during evolution. FLCN-1 and human folliculin share 23.4% identity and 39.9% similarity overall, and 21.8% identity and 37.0% similarity with their DENN domains. The residues absent in *flcn-1(ok975)* mutants are highlighted in red.

**Fig S3. In *rab-35(b1013)* mutants, recruitment of lysosomes to phagosomes is normal.**

(A) Time-lapse images monitoring the recruitment and fusion of lysosomes to the phagosomal surface (white arrows) after a phagosome forms (the “0 min” time point). Lysosomal fusion is monitored using a mcherry-tagged lysosomal lumen marker [NUC-1::mcherry]. The GFP-tagged PH domain of human phospholipase Cγ [PH(hPLCγ)::GFP], which labels PtdIns(4,5)P_2_ and extending phagosomes, is used to indicate the “0 min” time point when a phagosome forms. (B) Histogram displaying the range of time it takes for lysosomes to be recruited to the phagosomal surface in wild-type and *rab-35(b1013)* embryos. Phagosomes bearing cell corpses C1, C2, and C3 were scored. This time interval is defined as that between the “0 min” time point when a nascent phagosome is initially formed and the time point when a continuous CTNS-1::mRFP signal is observed on the phagosome. For each genotype, at least 15 phagosomes were scored. (C) Histogram displaying range of time it takes for lysosomes to fuse to phagosomes in wild-type and *rab-35(b1013)* embryos. Phagosomes bearing cell corpses C1, C2, and C3 were scored. The time span of lysosome fusion is measured as the time interval between “0 min” and the time point when the NUC-1::mCherry signal completely fills the phagosomal lumen. For each genotype, at least 15 phagosomes were scored.

**Fig S4. PIKI-1 recruitment to the phagosome is normal in *rab-35(b1013)* mutants.**

(A) Recruitment of the GFP-tagged class II PtdIns(3)P kinase GFP::PIKI-1 to nascent phagosomes was measured using live imaging of C1, C2, and C3 in wild-type and *rab-35(b1013)* mutant embryos at the 1.5-fold stage. The presence or absence of PIKI-1 on the phagosomes was scored on each phagosome and reported as a percentage for each genetic background. There was no significant decrease in the frequency of PIKI-1 recruitment in *rab-35(b1013)* mutants. (B) The intensity of PIKI-1 recruitment was measured in C1, C2, and C3. For each phagosome, the intensity of PIKI-1 signal was measured on the phagosome membrane and in the surrounding cytoplasm at the time point of maximal PIKI-1 phagosomal signal and expressed as a ratio. No statistically significant increase in PIKI-1 phagosomal intensity was observed in *rab-35(b1013)* mutants.

**Fig S5. *rab-35(b1013)* mutants display defects in apoptotic cell corpse recognition, but are normal for pseudopod extension.**

(A) The time it takes for engulfing cells to recognize cell corpses C1, C2, or C3 was determined in embryos of different genotypes using the GFP::CED-1ΔC reporter. The moment of recognition is defined as the first time point GFP is seen enriched in a region in contact between the engulfing and dying cell, with the moment of ventral enclosure used as a reference point (“0 min”). Histograms (a-d) and the summary (e-f) statistics are presented. For each strain, at least 15 engulfment events were scored. (B) The time it takes for C1, C2, or C3 to be internalized after they are recognized by the engulfing cells was determined in embryos of different genotypes using the GFP::CED-1ΔC reporter. Internalization is defined as the time interval between recognition of the dying cell by engulfing cells (“0 min”) and the time point that the nascent phagosome is formed. Histograms (a-d) and the summary (e-f) statistics are presented. For each strain, at least 15 engulfment events were scored.

**Fig S6. *rab-35(b1013)* mutants display an enhanced Ced phenotype in response to heat shock treatment.**

The mean numbers of apoptotic cell corpses were scored in 1.5-fold stage wild-type or *rab-35(b1013)* mutant embryos. The scoring was performed either with or without heat shock as well as in the presence or absence of transgenes overexpressing dominant negative GFP::RAB-5(S33N) under a heat shock promoter. For each data point, at least 15 animals were scored. Error bars indicate sd.

## References

1. Lockshin RA, Williams CM. Programmed cell death—II. Endocrine potentiation of the breakdown of the intersegmental muscles of silkmoths. J Insect Physiol. 1964 Aug 1;10(4):643–9.

2. Kerr JFR, Wyllie AH, Currie AR. Apoptosis: A Basic Biological Phenomenon with Wide-ranging Implications in Tissue Kinetics. Br J Cancer. 1972 Aug;26(4):239–57.

3. Elliott MR, Ravichandran KS. Clearance of apoptotic cells: implications in health and disease. J Cell Biol. 2010 Jun 28;189(7):1059–70.

4. Nagata S. Apoptosis and Clearance of Apoptotic Cells. Annu Rev Immunol [Internet]. 2018 [cited 2018 Apr 3];36(1). Available from: https://doi.org/10.1146/annurev-immunol-042617-053010

5. Sulston JE, Schierenberg E, White JG, Thomson JN. The embryonic cell lineage of the nematode Caenorhabditis elegans. Dev Biol. 1983 Nov 1;100(1):64–119.

6. Sulston JE, Horvitz HR. Post-embryonic cell lineages of the nematode, Caenorhabditis elegans. Dev Biol. 1977 Mar 1;56(1):110–56.

7. Gumienny TL, Lambie E, Hartwieg E, Horvitz HR, Hengartner MO. Genetic control of programmed cell death in the Caenorhabditis elegans hermaphrodite germline. Development. 1999 Mar 1;126(5):1011–22.

8. Yu X, Odera S, Chuang C-H, Lu N, Zhou Z. C. elegans Dynamin Mediates the Signaling of Phagocytic Receptor CED-1 for the Engulfment and Degradation of Apoptotic Cells. Dev Cell. 2006 Jun;10(6):743–57.

9. Zhou Z, Mangahas PM, Yu X. The Genetics of Hiding the Corpse: Engulfment and Degradation of Apoptotic Cells in C. elegans and D. melanogaster. In: Current Topics in Developmental Biology [Internet]. Academic Press; 2004 [cited 2017 Dec 1]. p. 91–143. Available from: http://www.sciencedirect.com/science/article/pii/S0070215304630043

10. Conradt B, Wu Y-C, Xue D. Programmed Cell Death During Caenorhabditis elegans Development. Genetics. 2016;203(4):1533–62.

11. Stenmark H, Olkkonen VM. The Rab GTPase family. Genome Biol. 2001;2(5):reviews3007.1-reviews3007.7.

12. Szatmári Z, Sass M. The autophagic roles of Rab small GTPases and their upstream regulators: a review. Autophagy. 2014 Jul;10(7):1154–66.

13. Lu N, Zhou Z. Membrane Trafficking and Phagosome Maturation During the Clearance of Apoptotic Cells. Int Rev Cell Mol Biol. 2012;293:269–309.

14. Gutierrez MG. Functional role(s) of phagosomal Rab GTPases. Small GTPases. 2013 Sep;4(3):148–58.

15. Kinchen JM, Doukoumetzidis K, Almendinger J, Stergiou L, Tosello-Trampont A, Sifri CD et al. A pathway for phagosome maturation during engulfment of apoptotic cells. Nat Cell Biol. 2008 May;10(5):556.

16. Lu N, Shen Q, Mahoney TR, Neukomm LJ, Wang Y, Zhou Z. Two PI 3-kinases and one PI 3-phosphatase together establish the cyclic waves of phagosomal PtdIns(3)P critical for the degradation of apoptotic cells. PLoS Biol. 2012 Jan;10(1):e1001245.

17. Mallo GV, Espina M, Smith AC, Terebiznik MR, Alemán A, Finlay BB et al. SopB promotes phosphatidylinositol 3-phosphate formation on Salmonella vacuoles by recruiting Rab5 and Vps34. J Cell Biol. 2008 Aug 25;182(4):741–52.

18. Yu X, Lu N, Zhou Z. Phagocytic Receptor CED-1 Initiates a Signaling Pathway for Degrading Engulfed Apoptotic Cells. PLoS Biol [Internet]. 2008 Mar [cited 2017 Nov 22];6(3). Available from: https://www.ncbi.nlm.nih.gov/pmc/articles/PMC2267821/

19. Harrison RE, Bucci C, Vieira OV, Schroer TA, Grinstein S. Phagosomes Fuse with Late Endosomes and/or Lysosomes by Extension of Membrane Protrusions along Microtubules: Role of Rab7 and RILP. Mol Cell Biol. 2003 Sep 15;23(18):6494–506.

20. He B, Yu X, Margolis M, Liu X, Leng X, Etzion Y, et al. Live-Cell Imaging in Caenorhabditis elegans Reveals the Distinct Roles of Dynamin Self-Assembly and Guanosine Triphosphate Hydrolysis in the Removal of Apoptotic Cells. Mol Biol Cell. 2010 Feb 15;21(4):610–29.

21. Lu Q, Zhang Y, Hu T, Guo P, Li W, Wang X. C. elegans Rab GTPase 2 is required for the degradation of apoptotic cells. Development. 2008 Mar 15;135(6):1069–80.

22. Mangahas PM, Yu X, Miller KG, Zhou Z. The small GTPase Rab2 functions in the removal of apoptotic cells in Caenorhabditis elegans. J Cell Biol. 2008 Jan 28;180(2):357–73.

23. Guo P, Hu T, Zhang J, Jiang S, Wang X. Sequential action of Caenorhabditis elegans Rab GTPases regulates phagolysosome formation during apoptotic cell degradation. Proc Natl Acad Sci. 2010 Oct 19;107(42):18016–21.

24. Levin R, Grinstein S, Canton J. The life cycle of phagosomes: formation, maturation, and resolution. Immunol Rev. 2016;273(1):156–79.

25. Cheng S, Wang K, Zou W, Miao R, Huang Y, Wang H, et al. PtdIns(4,5)P2 and PtdIns3P coordinate to regulate phagosomal sealing for apoptotic cell clearance. J Cell Biol. 2015 Aug 3;210(3):485–502.

26. Zhou Z, Hartwieg E, Horvitz HR. CED-1 Is a Transmembrane Receptor that Mediates Cell Corpse Engulfment in C. elegans. Cell. 2001 Jan 12;104(1):43–56.

27. Venegas V, Zhou Z. Two Alternative Mechanisms That Regulate the Presentation of Apoptotic Cell Engulfment Signal in Caenorhabditis elegans. Mol Biol Cell. 2007 Aug;18(8):3180–92.

28. Shen Q, He B, Lu N, Conradt B, Grant BD, Zhou Z. Phagocytic receptor signaling regulates clathrin and epsin-mediated cytoskeletal remodeling during apoptotic cell engulfment in C. elegans. Development. 2013 Aug 1;140(15):3230–43.

29. Liu QA, Hengartner MO. Candidate adaptor protein CED-6 promotes the engulfment of apoptotic cells in C. elegans. Cell. 1998 Jun 12;93(6):961–72.

30. Wu YC, Horvitz HR. The C. elegans cell corpse engulfment gene ced-7 encodes a protein similar to ABC transporters. Cell. 1998 Jun 12;93(6):951–60.

31. Gumienny TL, Brugnera E, Tosello-Trampont A-C, Kinchen JM, Haney LB, Nishiwaki K, et al. CED-12/ELMO, a Novel Member of the CrkII/Dock180/Rac Pathway, Is Required for Phagocytosis and Cell Migration. Cell. 2001 Oct 5;107(1):27–41.

32. Kang Y, Xu J, Liu Y, Sun J, Sun D, Hu Y, et al. Crystal structure of the cell corpse engulfment protein CED-2 in Caenorhabditis elegans. Biochem Biophys Res Commun. 2011 Jul 1;410(2):189–94.

33. Reddien PW, Horvitz HR. CED-2/CrkII and CED-10/Rac control phagocytosis and cell migration in Caenorhabditis elegans. Nat Cell Biol. 2000 Mar;2(3):131–6.

34. Chen W, Chen S, Yap SF, Lim L. The Caenorhabditis elegans p21-activated kinase (CePAK) colocalizes with CeRac1 and CDC42Ce at hypodermal cell boundaries during embryo elongation. J Biol Chem. 1996 Oct 18;271(42):26362–8.

35. Wu YC, Tsai MC, Cheng LC, Chou CJ, Weng NY. C. elegans CED-12 acts in the conserved crkII/DOCK180/Rac pathway to control cell migration and cell corpse engulfment. Dev Cell. 2001 Oct;1(4):491–502.

36. Mangahas PM, Zhou Z. Clearance of apoptotic cells in Caenorhabditis elegans. Semin Cell Dev Biol. 2005 Apr 1;16(2):295–306.

37. Hsu T-Y, Wu Y-C. Engulfment of Apoptotic Cells in C. elegans Is Mediated by Integrin α/SRC Signaling. Curr Biol. 2010 Mar 23;20(6):477–86.

38. Hsieh H-H, Hsu T-Y, Jiang H-S, Wu Y-C. Integrin α PAT-2/CDC-42 Signaling Is Required for Muscle-Mediated Clearance of Apoptotic Cells in Caenorhabditis elegans. PLOS Genet. 2012 May 17;8(5):e1002663.

39. WormBase11: Nematode Information Resource [Internet]. [cited 2017 Dec 6]. Available from: http://www.wormbase.org/#012-34-5

40. Gallegos ME, Balakrishnan S, Chandramouli P, Arora S, Azameera A, Babushekar A, et al. The C. elegans Rab Family: Identification, Classification and Toolkit Construction. PLOS ONE. 2012 Nov 21;7(11):e49387.

41. Sato M, Sato K, Liou W, Pant S, Harada A, Grant BD. Regulation of endocytic recycling by C. elegans Rab35 and its regulator RME-4, a coated-pit protein. EMBO J. 2008 Apr 23;27(8):1183–96.

42. Rompay LV, Borghgraef C, Beets I, Caers J, Temmerman L. New genetic regulators question relevance of abundant yolk protein production in C. elegans. Sci Rep [Internet]. 2015 Nov 10 [cited 2018 Mar 28];5. Available from: https://www.ncbi.nlm.nih.gov/pmc/articles/PMC4639837/

43. Lu N, Yu X, He X, Zhou Z. Detecting apoptotic cells and monitoring their clearance in the nematode Caenorhabditis elegans. Methods Mol Biol Clifton NJ. 2009;559:357–70.

44. Chaineau M, Ioannou MS, McPherson PS. Rab35: GEFs, GAPs and Effectors. Traffic. 2013 Nov 1;14(11):1109–17.

45. Liu B, Du H, Rutkowski R, Gartner A, Wang X. LAAT-1 is the Lysosomal Lysine/Arginine Transporter that Maintains Amino Acid Homeostasis. Science. 2012 Jul 20;337(6092):351–4.

46. Lu N, Shen Q, Mahoney TR, Liu X, Zhou Z. Three sorting nexins drive the degradation of apoptotic cells in response to PtdIns(3)P signaling. Mol Biol Cell. 2011 Feb 1;22(3):354–74.

47. Ellis HM, Horvitz HR. Genetic control of programmed cell death in the nematode C. elegans. Cell. 1986 Mar 28;44(6):817–29.

48. Reddien PW, Horvitz HR. The engulfment process of programmed cell death in caenorhabditis elegans. Annu Rev Cell Dev Biol. 2004;20:193–221.

49. Hsu C, Morohashi Y, Yoshimura S, Manrique-Hoyos N, Jung S, Lauterbach MA et al. Regulation of exosome secretion by Rab35 and its GTPase-activating proteins TBC1D10A– C. J Cell Biol. 2010 Apr 19;189(2):223–32.

50. Biesemann A, Gorontzi A, Barr F, Gerke V. Rab35 protein regulates evoked exocytosis of endothelial Weibel-Palade bodies. J Biol Chem. 2017 14;292(28):11631–40.

51. Patino-Lopez G, Dong X, Ben-Aissa K, Bernot KM, Itoh T, Fukuda M, et al. Rab35 and Its GAP EPI64C in T Cells Regulate Receptor Recycling and Immunological Synapse Formation. J Biol Chem. 2008 Jun 27;283(26):18323–30.

52. Walseng E, Bakke O, Roche PA. Major Histocompatibility Complex Class II-Peptide Complexes Internalize Using a Clathrin- and Dynamin-independent Endocytosis Pathway. J Biol Chem. 2008 May 23;283(21):14717–27.

53. Gao Y, Balut CM, Bailey MA, Patino-Lopez G, Shaw S, Devor DC. Recycling of the Ca2+-activated K+ Channel, KCa2.3, Is Dependent upon RME-1, Rab35/EPI64C, and an N-terminal Domain. J Biol Chem. 2010 Jun 4;285(23):17938–53.

54. Sheehan P, Zhu M, Beskow A, Vollmer C, Waites CL. Activity-Dependent Degradation of Synaptic Vesicle Proteins Requires Rab35 and the ESCRT Pathway. J Neurosci Off J Soc Neurosci. 2016 17;36(33):8668–86.

55. Kouranti I, Sachse M, Arouche N, Goud B, Echard A. Rab35 Regulates an Endocytic Recycling Pathway Essential for the Terminal Steps of Cytokinesis. Curr Biol. 2006 Sep 5;16(17):1719–25.

56. Chesneau L, Dambournet D, Machicoane M, Kouranti I, Fukuda M, Goud B, et al. An ARF6/Rab35 GTPase Cascade for Endocytic Recycling and Successful Cytokinesis. Curr Biol. 2012 Jan 24;22(2):147–53.

57. Shim J, Lee S-M, Lee MS, Yoon J, Kweon H-S, Kim Y-J. Rab35 mediates transport of Cdc42 and Rac1 to the plasma membrane during phagocytosis. Mol Cell Biol. 2010 Mar;30(6):1421–33.

58. Dambournet D, Machicoane M, Chesneau L, Sachse M, Rocancourt M, Marjou AE et al. Rab35 GTPase and OCRL phosphatase remodel lipids and F-actin for successful cytokinesis. Nat Cell Biol. 2011 Aug;13(8):981.

59. Chevallier Julien, Koop Charles, Srivastava Archana, Petrie Ryan J., Lamarche-Vane Nathalie, Presley John F. Rab35 regulates neurite outgrowth and cell shape. FEBS Lett. 2009 Mar 14;583(7):1096–101.

60. Marat AL, McPherson PS. The Connecdenn Family, Rab35 Guanine Nucleotide Exchange Factors Interfacing with the Clathrin Machinery. J Biol Chem. 2010 Apr 2;285(14):10627–37.

61. Zhang J, Fonovic M, Suyama K, Bogyo M, Scott MP. Rab35 controls actin bundling by recruiting fascin as an effector protein. Science. 2009 Sep 4;325(5945):1250–4.

62. Minowa-Nozawa A, Nozawa T, Okamoto-Furuta K, Kohda H, Nakagawa I. Rab35 GTPase recruits NDP52 to autophagy targets. EMBO J. 2017 15;36(18):2790–807.

63. Wheeler DB, Zoncu R, Root DE, Sabatini DM, Sawyers CL. Identification of an oncogenic RAB protein. Science. 2015 Oct 9;350(6257):211–7.

64. Egami Y, Fukuda M, Araki N. Rab35 regulates phagosome formation through recruitment of ACAP2 in macrophages during FcγR-mediated phagocytosis. J Cell Sci. 2011 Nov 1;124(21):3557–67.

65. Egami Y, Fujii M, Kawai K, Ishikawa Y, Fukuda M, Araki N. Activation-Inactivation Cycling of Rab35 and ARF6 Is Required for Phagocytosis of Zymosan in RAW264 Macrophages [Internet]. Journal of Immunology Research. 2015 [cited 2017 Dec 6]. Available from: https://www.hindawi.com/journals/jir/2015/429439/

66. Stroschein-Stevenson SL, Foley E, O’Farrell PH, Johnson AD. Identification of Drosophila gene products required for phagocytosis of Candida albicans. PLoS Biol. 2006 Jan;4(1):e4.

67. Smith AC, Heo WD, Braun V, Jiang X, Macrae C, Casanova JE et al. A network of Rab GTPases controls phagosome maturation and is modulated by Salmonella enterica serovar Typhimurium. J Cell Biol. 2007 Jan 29;176(3):263–8.

68. Cai H, Reinisch K, Ferro-Novick S. Coats, tethers, Rabs, and SNAREs work together to mediate the intracellular destination of a transport vesicle. Dev Cell. 2007 May;12(5):671–82.

69. Kinchen JM, Ravichandran KS. Identification of two evolutionarily conserved genes regulating processing of engulfed apoptotic cells. Nature. 2010 Apr 1;464(7289):778–82.

70. Li W, Zou W, Zhao D, Yan J, Zhu Z, Lu J, et al. C. elegans Rab GTPase activating protein TBC-2 promotes cell corpse degradation by regulating the small GTPase RAB-5. Development. 2009 Jul 15;136(14):2445–55.

71. Klinkert K, Echard A. Rab35 GTPase: A Central Regulator of Phosphoinositides and F-actin in Endocytic Recycling and Beyond. Traffic. 2016 Oct 1;17(10):1063–77.

72. Nookala RK, Langemeyer L, Pacitto A, Ochoa-Montaño B, Donaldson JC, Blaszczyk BK et al. Crystal structure of folliculin reveals a hidDENN function in genetically inherited renal cancer. Open Biol. 2012 Aug 1;2(8):120071.

73. Zheng J, Duan B, Sun S, Cui J, Du J, Zhang Y. Folliculin Interacts with Rab35 to Regulate EGF-Induced EGFR Degradation. Front Pharmacol. 2017;8:688.

74. Pfeffer S, Aivazian D. Targeting Rab GTPases to distinct membrane compartments. Nat Rev Mol Cell Biol. 2004 Nov;5(11):886–96.

75. Heo WD, Inoue T, Park WS, Kim ML, Park BO, Wandless TJ et al. PI(3,4,5)P3 and PI(4,5)P2 lipids target proteins with polybasic clusters to the plasma membrane. Science. 2006 Dec 1;314(5804):1458–61.

76. Gavriljuk K, Itzen A, Goody RS, Gerwert K, Kötting C. Membrane extraction of Rab proteins by GDP dissociation inhibitor characterized using attenuated total reflection infrared spectroscopy. Proc Natl Acad Sci. 2013 Aug 13;110(33):13380–5.

77. Li F, Yi L, Zhao L, Itzen A, Goody RS, Wu Y-W. The role of the hypervariable C-terminal domain in Rab GTPases membrane targeting. Proc Natl Acad Sci. 2014 Feb 18;111(7):2572–7.

78. Kwiatkowska K. One lipid, multiple functions: how various pools of PI(4,5)P(2) are created in the plasma membrane. Cell Mol Life Sci CMLS. 2010 Dec;67(23):3927–46.

79. Ellis RE, Jacobson DM, Horvitz HR. Genes required for the engulfment of cell corpses during programmed cell death in Caenorhabditis elegans. Genetics. 1991 Sep 1;129(1):79–94.

80. D’mello V, Birge RB. Apoptosis: Conserved Roles for Integrins in Clearance. Curr Biol. 2010 Apr 13;20(7):R324–7.

81. Neukomm LJ, Zeng S, Frei AP, Huegli PA, Hengartner MO. Small GTPase CDC-42 promotes apoptotic cell corpse clearance in response to PAT-2 and CED-1 in C. elegans. Cell Death Differ. 2014 Jun;21(6):845.

82. Carlson JM, Doyle J. Complexity and robustness. Proc Natl Acad Sci U S A. 2002 Feb 19;99 Suppl 1:2538–45.

83. Kitano H. Cancer robustness: tumour tactics. Nature. 2003 Nov 13;426(6963):125.

84. Kitano H. Biological robustness. Nat Rev Genet. 2004 Nov;5(11):826–37.

85. Rovenský J, Tuchynová A. Systemic lupus erythematosus in the elderly. Autoimmun Rev. 2008 Jan;7(3):235–9.

86. Thakkar JP, McCarthy BJ, Villano JL. Age-specific cancer incidence rates increase through the oldest age groups. Am J Med Sci. 2014 Jul;348(1):65–70.

87. Erwig L-P, Henson PM. Immunological Consequences of Apoptotic Cell Phagocytosis. Am J Pathol. 2007 Jul 1;171(1):2–8.

88. Brenner S. The genetics of Caenorhabditis elegans. Genetics. 1974 May;77(1):71–94.

89. Jin Y. C elegans: A Practical Approach. Hope I, editor. New York: Oxford University Press; 1999. 69–96 p. (Transformation).

90. Bloom L, Horvitz HR. The Caenorhabditis elegans gene unc-76 and its human homologs define a new gene family involved in axonal outgrowth and fasciculation. Proc Natl Acad Sci U S A. 1997 Apr 1;94(7):3414–9.

91. Kamath RS, Martinez-Campos M, Zipperlen P, Fraser AG, Ahringer J. Effectiveness of specific RNA-mediated interference through ingested double-stranded RNA in Caenorhabditis elegans. Genome Biol. 2001;2(1):RESEARCH0002.

92. Kamath RS, Fraser AG, Dong Y, Poulin G, Durbin R, Gotta M, et al. Systematic functional analysis of the Caenorhabditis elegans genome using RNAi. Nature. 2003 Jan 16;421(6920):231–7.

93. Kamath RS, Ahringer J. Genome-wide RNAi screening in Caenorhabditis elegans. Methods San Diego Calif. 2003 Aug;30(4):313–21.

